# Engineered T cells controlled by granulocyte colony-stimulating factor eradicate murine melanoma without lymphodepletion or interleukin-2

**DOI:** 10.64898/2026.10.22.760485

**Authors:** Sam C. Preshaw, Megan E. Fuller, Cameron J. Powell, Katy Milne, Kassandra Sewell, Mitchell B. Adamson, Alexi Pearson-Lund, Marco Joly, Sydney Goodwin, Niloufar Shakerian, Natalie J.S. Taylor, Ming Zi Xu, Danielle Klassen, Taise Bryce, Julian Smazynski, Martin J. Boulanger, Brad H. Nelson

## Abstract

Tumour-infiltrating lymphocyte (TIL) therapy can be effective against metastatic melanoma, including immune checkpoint-resistant disease, but current protocols require lymphodepletion and high-dose interleukin-2 (IL-2), which can cause substantial toxicity. To overcome this limitation, we engineered a chimeric receptor, G2R, that generates an IL-2-like proliferative signal in response to granulocyte colony-stimulating factor (G-CSF), a well-tolerated cytokine widely used in oncology. Human T cells expressing G2R expanded in vitro in response to G-CSF in place of IL-2 or interleukin-15 (IL-15), and the three cytokines induced broadly similar intracellular signalling and cytokine secretion profiles. In immune-competent mice bearing established B16-F10 melanoma, adoptive transfer of G2R-expressing, tumour-reactive CD8^+^ T cells followed by systemic G-CSF treatment induced profound T cell expansion. This was accompanied by complete, durable tumour regression in over 75% of animals without the need for lymphodepletion, exogenous IL-2 or any other therapy. G-CSF treatment induced transient neutrophil expansion in peripheral blood but no increase in intratumoural neutrophils or macrophages. G2R-expressing memory T cells persisted long-term and could be re-expanded upon G-CSF re-treatment, enabling rejection of second tumour challenges. Thus, G-CSF can be repurposed to enable clinician-controlled anti-tumour T cell responses of profound magnitude, efficacy and durability, an approach that is readily translatable to humans.

## Introduction

Immune checkpoint blockade (ICB) has revolutionized the treatment of advanced metastatic melanoma, yet roughly half of patients still die of their disease, emphasizing the need for new treatment strategies^1–3^. A promising alternative called TIL therapy involves surgical resection of a tumour sample and ex vivo expansion of tumour-associated T cells with IL-2 followed by a rapid expansion protocol^4–6^. Patients receive lymphodepleting chemotherapy followed by a single dose of 1 x 10⁹ – 2 × 10¹¹ TILs and up to six cycles of high-dose IL-2. In pooled analysis, current protocols achieve objective and complete response rates of 34% and 10%, respectively^7^. However, lymphodepletion and, in particular, high-dose IL-2 frequently cause significant toxicities, including fever, hypotension, pulmonary oedema, and neurotoxicity, such that many patients receive fewer than the recommended number of IL-2 cycles^8,9^. In theory, lymphodepletion could also impair the efficacy of TIL therapy given accumulating evidence that host T cells and natural killer (NK) cells can make important contributions to the anti-tumour activity of chimeric antigen receptor (CAR)-T cells and other T cell therapies^10,11^.

A wide variety of strategies have been investigated to improve the efficacy and safety of TIL therapy. Several groups have engineered mutated variants of IL-2 that exhibit reduced binding to IL-2 receptor alpha (IL-2Rα), thereby favouring the expansion of CD8^+^ T cells and NK cells over regulatory T cells (Tregs)^12–14^. In clinical trials, such IL-2 variants have indeed reduced Treg expansion and some IL-2-related toxicities while promoting expansion of CD8^+^ T cells and NK cells; however, only modest anti-tumour responses have been reported^15^. In another strategy, IL-2 was fused to an IL-21 receptor alpha (IL-21Rα) binding element, creating a hybrid cytokine that induces a combined IL-2 and IL-21 signal^16^. While achieving moderate efficacy in mice bearing B16-F10 melanoma, this came at the expense of considerable body weight loss, which is indicative of immune-related toxicity. “Switch receptors” have been designed that bind immunosuppressive cytokines such as transforming growth factor beta (TGF-β), IL-4, or IL-10 yet generate immunostimulatory signals resembling those induced by IL-2, IL-7 or IL-21^17–19^. Although these strategies have shown moderate efficacy in CAR-T cells, their activity is not externally tunable and relies on endogenous cytokines whose levels may vary substantially within and between patients. Yet another approach has been to engineer CAR-T cells or TIL to express constitutively active IL-2 or IL-7 receptors, which can eliminate the need for exogenous cytokine^20–22^. Clinical experience with GD2-directed CAR-T cells paired with constitutively active IL-7 receptor has demonstrated partial responses and manageable levels of toxicity, yet it remains to be seen whether such a strategy can induce long-term tumour regressions^23^.

Two additional engineering strategies have been developed to confer more precise control over TIL products in vivo. In one case, complementary variants of IL-2 and the IL-2 receptor β (IL-2Rβ) subunit were engineered to create an orthogonal system wherein “ortho-IL-2” is used to selectively stimulate T cells expressing “ortho-IL-2Rβ”. While this approach achieved enhanced survival in the B16-F10 model, complete regression (CR) of tumours was not observed^24^. Furthermore, dose-dependent toxicities were seen in a CD19-directed CAR T cell leukaemia model, raising concerns about the safety of combining this cytokine:receptor system with CAR-T cells^25,26^. Variations of this system have been designed that emulate signals other than IL-2, including IL-9, IL-21, IL-22, and IL-23^27,28^. Several of these designs have demonstrated strong efficacy, yet all still require lymphodepletion and some have shown evidence of toxicity. In the second approach, TIL were engineered to express a membrane-bound form of IL-15 (mbIL15) whose expression is controlled by acetazolamide, an FDA-approved small molecule drug^29^. This strategy showed strong efficacy in a patient-derived xenograft model of melanoma TIL therapy and is yielding encouraging preliminary results in phase I/II clinical trials for the treatment of ICB-resistant advanced melanoma when combined with low-dose lymphodepletion^30^.

As an alternative to the above strategies, we investigated whether an IL-2-like signal could be delivered to T cells using G-CSF, a cytokine which is routinely and safely given to cancer patients (including those receiving TIL therapy) to manage neutropenia^9,31^. We designed a chimeric receptor, G2R, which fuses the extracellular, transmembrane, and Janus-kinase (JAK) binding domains of the G-CSF receptor to key signalling domains from IL-2Rβ (Fig. 1a). When expressed in T cells, G2R closely emulated IL-2 signalling and induced profound T cell expansion and anti-tumour activity in vivo, eliminating the need for lymphodepletion, IL-2 or any other therapy in the challenging B16-F10 murine melanoma model. Moreover, G2R-expressing memory T cells could be re-expanded to therapeutically effective levels by re-treatment with G-CSF, without the need for any other therapies. As expected, G-CSF also induced neutrophil expansion, but this was well-tolerated, reversible, and did not impair anti-tumour efficacy. Thus, G2R technology provides a clinically feasible strategy to improve both the safety and efficacy of TIL therapy and potentially other cell therapies.

**Figure 1.**
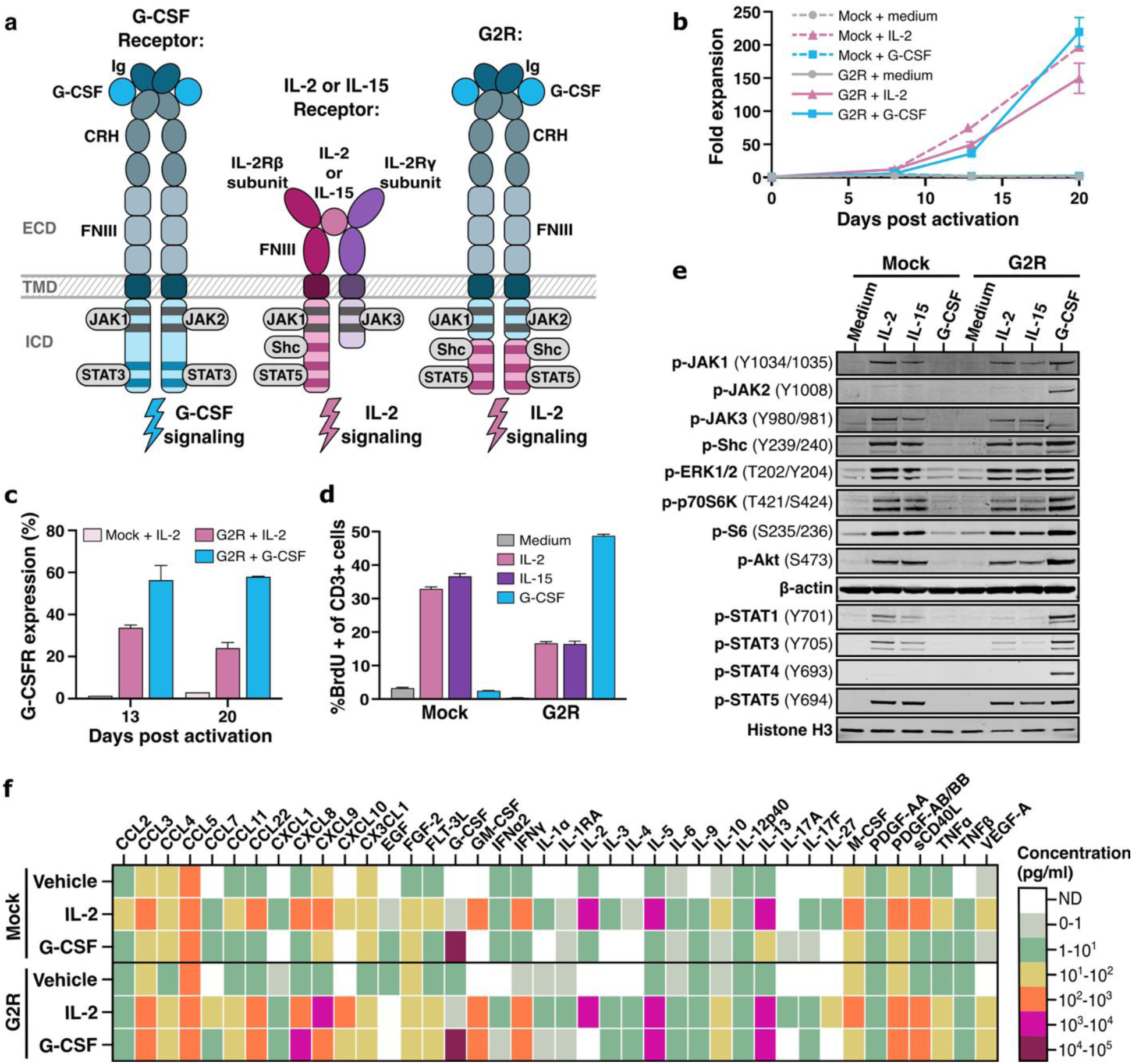
G2R closely emulates IL-2R signalling and drives human T cell expansion and cytokine production in vitro. **(a)** Structure of wild-type (WT) G-CSFR, IL-2R, and G2R, including the extracellular domain (ECD), transmembrane domain (TMD), and intracellular domain (ICD) of each receptor. Also shown are the immunoglobulin-like (Ig), cytokine receptor-homologous (CRH), and fibronectin type III (FNIII) domains of the G-CSFR and IL-2R ECDs, and the JAK, Shc and STAT binding sites of the G-CSFR and IL-2Rβ ICDs. **(b-f)** Human CD4^+^ and CD8^+^ T cells from peripheral blood were transduced to express G2R and assessed in vitro for cytokine responsiveness relative to mock-transduced controls. **(b)** T cell expansion in the presence of G-CSF, IL-2 or media. **(c)** G2R surface expression on days 13 and 20 of expansion as measured by flow cytometry for G-CSFR. **(d)** BrdU-incorporation assay using mock-transduced or G2R-transduced T cells expanded for 13 days in IL-2 or G-CSF, respectively. T cells were washed and then re-stimulated for 2 days with IL-2 (300 IU/mL), IL-15 (10 ng/mL), G-CSF (100 ng/mL), or media. **(e)** Western blots showing phosphorylation (p) of the indicated proteins in T cells stimulated for 20 minutes with the indicated cytokines. **(f)** Heat map showing the levels of the indicated cytokines in supernatants from T cell cultures stimulated for 4 days with vehicle, IL-2 or G-CSF. Not shown are cytokines that were below the limit of detection (IL-7, IL-17E, IL-22) or with values below 1 pg/mL (IL-1β, IL-12p70, IL-15, IL-18, TGFα). Data in panels b, c and d are presented as means +/-SD of two technical replicates. Data in panel f are presented as means of two technical replicates.

## Results

### A chimeric G-CSF/IL-2 receptor (“G2R”) emulates IL-2 signalling in human T cells

To enable the delivery of an IL-2-like signal to T cells in response to G-CSF, we designed a chimeric receptor, G2R, composed of the extracellular, transmembrane and JAK1 and JAK2 binding domains of the G-CSF receptor fused to the C-terminal portion of IL-2Rβ containing the src homology 2 domain-containing (Shc)- and signal transducer and activator of transcription (STAT) 5-binding sites (Fig. 1a, Table S1). G2R was introduced by lentiviral transduction into primary human CD4^+^ and CD8^+^ T cells, which were then cultured for 20 days in media, IL-2, or G-CSF. As expected, mock-transduced T cells expanded in response to IL-2 (197.0-fold) but not G-CSF. In contrast, G2R⁺ T cells expanded to similar levels in response to either G-CSF or IL-2 (219.4-versus 149.7-fold expansion, respectively) and maintained similarly high viability in the two cytokines (93.0% and 95.2%, respectively) compared to media alone (61.9%) (Fig. 1b, Fig. S1a). G2R was expressed on 58.0% versus 24.0% of G2R-transduced T cells expanded in G-CSF or IL-2, respectively (Fig. 1c), indicating that G-CSF positively selected for G2R expression. G2R^+^ T cells that had been expanded in G-CSF remained dependent on exogenous cytokine, as demonstrated by subsequent proliferative responses to G-CSF, IL-2 or IL-15 but not media alone (Fig. 1d). The half-maximal effective concentration (EC₅₀) of G-CSF was 8.6 ng/mL (Fig. S1b), which is slightly greater than that reported for the native G-CSFR in other cell types^32,33^ and well above physiological serum levels of G-CSF (15-22 pg/mL) in humans^34–36^. Thus, G2R confers G-CSF-dependent control of T cell proliferation and viability, completely replacing the need for IL-2 or IL-15.

To assess intracellular signalling events induced by G2R compared to the native IL-2R, we stimulated G2R-expressing T cells with G-CSF, IL-2, or IL-15, and analysed phosphorylation of key intracellular signalling proteins by western blot. Mock-transduced T cells served as a negative control. As expected, IL-2 and IL-15 induced tyrosine phosphorylation of JAK1 and JAK3 in both mock-transduced and G2R-expressing T cells^37,38^. In G2R-expressing T cells, G-CSF induced tyrosine phosphorylation of JAK1 and JAK2, which was expected given that G2R uses the Box 1 and 2 regions of G-CSFR^39^. Downstream of the JAKs, G-CSF, IL-2, and IL-15 induced similar phosphorylation of Shc, extracellular signal-regulated kinase (ERK), protein kinase B (Akt), and ribosomal protein S6 kinase beta-1 (p70S6K), as well as the transcription factors STAT1, STAT3, and STAT5 (Fig. 1e). G-CSF also induced tyrosine phosphorylation of STAT4, which has been reported as an IL-2-induced signalling event in NK cells but not T cells^40^.

Flow cytometry of G2R-expressing T cells revealed that the frequency of cells containing tyrosine phosphorylated STAT5 (P-STAT5) was similar with G-CSF versus IL-2 stimulation (Fig. S1c). The median fluorescence intensity (MFI) of P-STAT5 peaked at 20 minutes for both G-CSF and IL-2, with G-CSF inducing an MFI 1.8-fold greater than that of IL-2 (Fig. S1d). As expected, we did not observe any signalling events in mock-transduced T cells stimulated with G-CSF. Therefore, apart from JAK2 and STAT4 phosphorylation, G2R faithfully emulates the intracellular signalling events induced by the native IL-2 and IL-15 receptors.

In addition to promoting T-cell proliferation and viability, IL-2 stimulates the secretion of diverse cytokines and chemokines^41,42^. To examine whether G2R induces similar effects, we stimulated day 12 cultures of G2R⁺ or mock-transduced T cells with G-CSF or IL-2, collected supernatants after four days of incubation, and performed an array-based analysis of 48 cytokines and chemokines. G-CSF and IL-2 induced similar patterns of cytokine/chemokine secretion, including chemokine ligand 3 (CCL3), chemokine ligand 9 (CXCL9), granulocyte-macrophage colony-stimulating factor (GM-CSF), interferon gamma (IFN-γ), IL-13, soluble cluster of differentiation 40 ligand (sCD40L) and tumour necrosis factor alpha (TNFα) (Fig. 1f).

### G-CSF induces profound expansion, persistence and anti-tumour activity of G2R-expressing tumour-specific T cells in vivo

Having validated the signalling fidelity of G2R in vitro, we next investigated whether G-CSF could be used to control the expansion and anti-tumour activity of G2R-expressing T cells in vivo. This was evaluated using the murine B16-F10 melanoma model and TCR-transgenic CD8^+^ T cells specific for the gp100 antigen (pmel-1 T cells). This is a well-established experimental model that typically requires the use of lymphodepletion (usually by total body irradiation), high-dose IL-2, and additional treatments such as vaccination to achieve significant anti-tumour responses^43–46^. Wild-type, fully immune-competent mice bearing established (day 7) subcutaneous B16-F10 tumours underwent adoptive cell transfer (ACT) with 1 x 10^7^ pmel-1 T cells expressing G2R. Mock-transduced pmel-1 T cells served as a negative control (Fig. 2a). We used a fully murine version of G2R to minimize immunogenicity and confirmed that it was expressed and signalled as expected in pmel-1 T cells (Fig. S2, a to c). Mice were randomized to receive G-CSF, IL-2 or vehicle for three weeks post-ACT. We used pegylated G-CSF (PEG-GCSF), as this formulation is commonly used in cancer patients and requires less frequent administration. As pegylated IL-2 is not readily available, we used non-pegylated IL-2 with daily dosing, a regimen that is commonly used in the B16-F10/pmel-1 model^47–49^. To enable a sufficient number of control groups and adequate sample sizes, three independent experiments were performed (Table S2), and data were combined where indicated.

**Figure 2.**
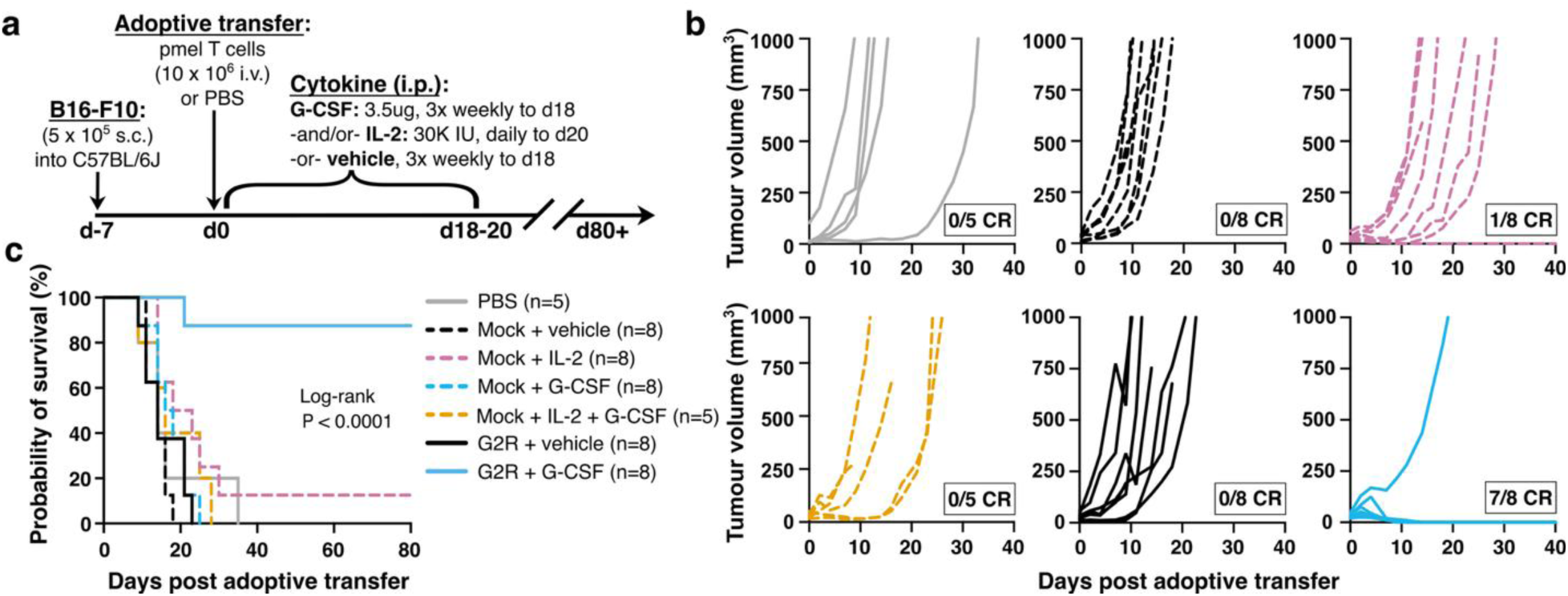
Treatment with G2R^+^ T cells + G-CSF induces complete, durable regression of established B16-F10 melanoma in immune-competent mice. **(a)** C57Bl/6J mice (n = 5 to 8 per group) were implanted subcutaneously with 5 x 10^5^ B16-F10 melanoma cells seven days prior to intravenous adoptive transfer of 1 x 10^7^ mock-transduced or G2R-transduced pmel-1 T cells. Mice were randomized to receive intraperitoneal injections of pegylated G-CSF (3x weekly to day 18), IL-2 (daily to day 20) or vehicle alone. **(b)** Tumour volumes as measured by caliper three times weekly. **(c)** Kaplan-Meier survival curves with all treatment groups compared by log-rank (Mantel-Cox) test.

In mice treated with mock-transduced pmel-1 T cells + vehicle or + G-CSF, no complete regressions (CR) were observed (0/12 CR for vehicle and 0/13 CR for G-CSF over two experiments) (Fig. 2, b and c, Fig. S2, d to g), demonstrating that unmodified pmel-1 T cells were ineffective under these conditions. Mice that received mock-transduced T cells + IL-2 showed a slight therapeutic benefit, with 1/13 mice exhibiting CR (7.7%, over two experiments). Of the mice that received G2R^+^ pmel-1 T cells + vehicle or + IL-2, none showed therapeutic benefit (0/13 CR for vehicle and 0/3 CR for IL-2, over two experiments). Strikingly, however, the combination of G2R^+^ T cells + G-CSF resulted in CR in 14/18 mice (77.8%, over three experiments). These responses were rapid, with tumours becoming unpalpable within 9 to 15 days of ACT. The responses were also durable, with none of the 14 mice that experienced CR demonstrating tumour recurrence over observation periods of 74 to 98 days (Fig. 2, b and c, Fig. S2, d to g).

To assess whether the remarkable efficacy seen with the G2R^+^ T cell + G-CSF condition could result from the combined effects of delivering an IL-2 signal to T cells and a G-CSF signal to myeloid cells, we included an additional control group consisting of mock-transduced T cells + IL-2 + G-CSF (Fig. 2, b and c). This combination resulted in a modest delay in tumour growth but 0/5 CRs, which was no better than the response seen with mock-transduced pmel-1 T cells + IL-2. Thus, the strong efficacy observed as a result of combined treatment with G2R^+^ T cells + G-CSF can be attributed to G2R-mediated signalling in pmel-1 T cells rather than systemic effects of G-CSF on myeloid or other cell types.

The number of circulating pmel-1 T cells was tracked over time by flow cytometry. Across all control groups, including mice receiving pmel-1 T cells + IL-2, minimal expansion and persistence of pmel-1 T cells was observed. For example, in the experiment shown in Fig. 3a, mean values of 2.0 x 10^4^ cells/mL and 2.0 x 10^3^ cells/mL were observed on days 4 and 9, respectively, in mice receiving mock-transduced T cells + IL-2. In contrast, in mice treated with G2R^+^ T cells + G-CSF, we observed rapid and profound expansion of pmel-1 T cells. Peak levels were reached by day 4 post ACT, when the level of pmel-1 T cells exceeded that of the vehicle-treated groups by anywhere from 13.4- to 184.4-fold (mean = 80.0-fold, n = 8 animals) and averaged 5.8 x 10^5^ cells/mL (29.0-fold higher than the mock-transduced T cells + IL-2 group), representing 62.2% of all circulating CD8^+^ T cells (Fig. S2h). We observed a positive correlation between the number of circulating pmel-1 T cells and the percentage exhibiting a T effector memory (Tem, CD44⁺CD62L^-^) phenotype on day 3 to 4 post ACT in the G2R^+^ pmel-1 T cells + G-CSF-treated group. This trend was consistent across the three studies and significant in one (P = 0.0008, n = 8 animals in Study 2) (Fig. S2i). By day 9, pmel-1 levels contracted to an average of 1.3 x 10^5^ cells/mL, which was 64.6-fold higher than the mock-transduced T cells + IL-2 group and 25.2- to 117.4-fold higher than all other control groups.

**Figure 3.**
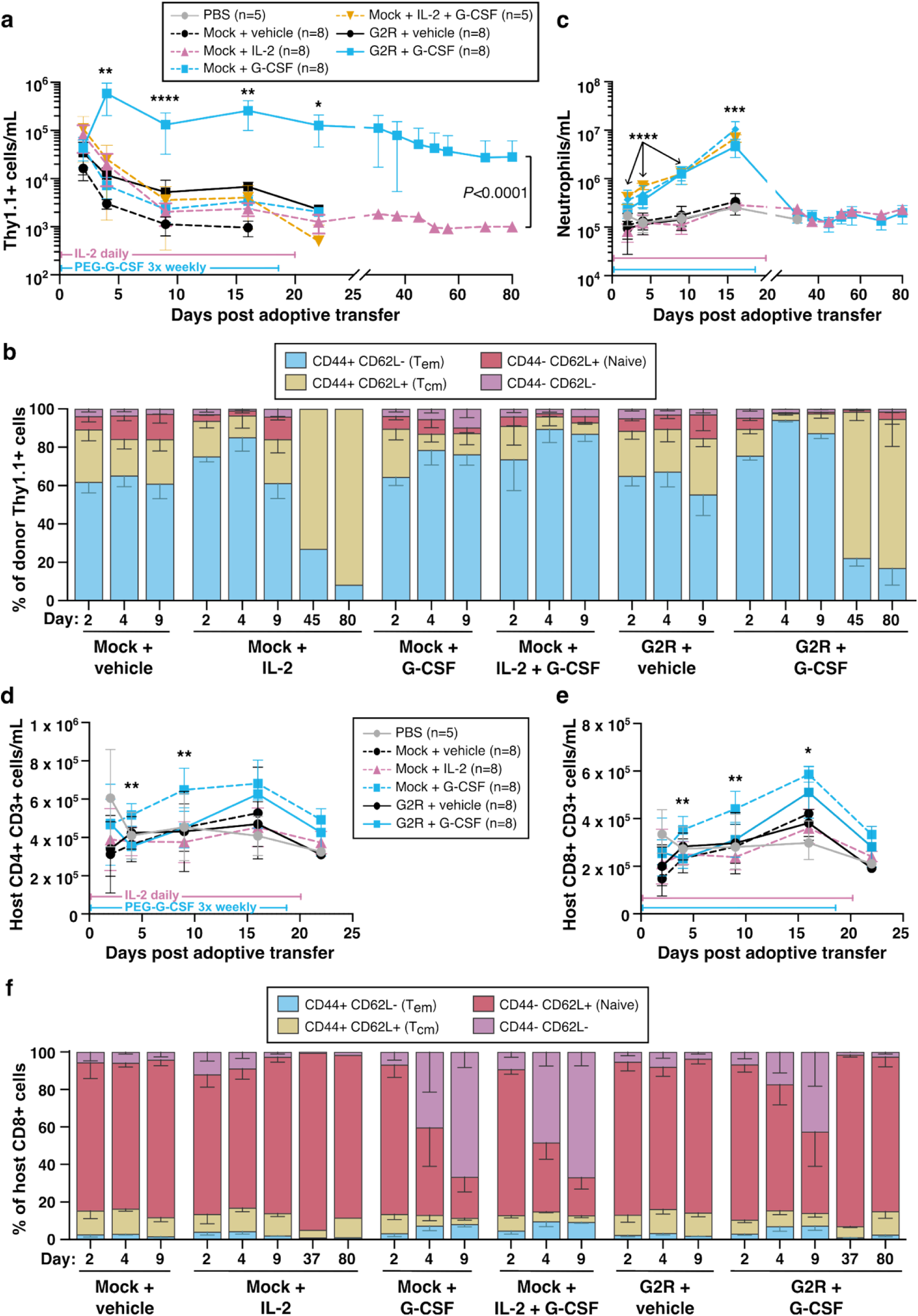
G-CSF induces profound expansion and persistence of G2R-expressing pmel-1 T cells and reversible neutrophil expansion in vivo. Serial peripheral blood samples were analysed by flow cytometry to measure: **(a)** Adoptively transferred Thy1.1^+^ pmel-1 T cells. Mann-Whitney test of mock + IL-2 vs. G2R + G-CSF: P < 0.0001. **(b)** Neutrophils. **(c)** Memory phenotypes of Thy1.1^+^ pmel-1 T cells based on CD44 and CD62L expression. Dunnett’s T3 multiple comparisons test was used to compare mean %Tem phenotype on day 4 for the G2R^+^ T cells + vehicle vs. G2R^+^ T cells + G-CSF groups (P = 0.0004) and the mock T cells + vehicle vs. G2R^+^ T cells + G-CSF groups (P <0.0001). **(d)** Host (Thy1.1^-^) CD4^+^CD3^+^ T cells. **(e)** Host (Thy1.1^-^) CD8^+^CD3^+^ T cells. **(f)** Memory phenotypes of host (Thy1.1^-^) CD8^+^CD3^+^ T cells based on CD44 and CD62L expression. Panels **a, c, d, e**: Welch’s ANOVA tests on days 2, 4, 9 and 16. *P < 0.05; **P < 0.01; ***P < 0.001; ****P < 0.0001. Data are presented as means +/-SD of 5 to 8 mice per group, as indicated. Animals were treated, as indicated, with PEG-GCSF 3x weekly until day 18, and IL-2 daily until day 20. Some group sizes decreased over time as mice reached the humane endpoint.

Across the three experiments, the 4/18 mice that failed to exhibit CR despite receiving G2R^+^ pmel-1 T cells + G-CSF had significantly fewer circulating pmel-1 T cells on day 3 to 4 after ACT than the 14/18 mice that underwent CR (mean = 8.2 x 10^4^ versus 1.2 x 10^6^ cells/mL, P = 0.02) (Fig. S2j). Thus, the success of this treatment appears to depend on obtaining a sufficiently high ratio of T cells to tumour cells early in the response to oppose the rapid growth of B16-F10 tumours.

In the 14/18 mice that experienced CRs, we assessed the persistence of G2R⁺ pmel-1 T cells after tumour clearance. On days 15 to 17, average pmel-1 T cell levels were 70.3-fold higher in these mice than in the 9/13 mice that were treated with G2R^+^ T cells + vehicle and were still alive at these timepoints (mean = 2.6 x 10^5^ cells/mL versus 3.7 x 10^3^ cells/mL, respectively; P = 0.0037) (Fig. S2k). Notably, however, within the CR group, individual pmel-1 values ranged from 2.8 x 10^2^ to 8.8 x 10^5^ cells/mL on days 15 to 17, a difference of more than 3,100-fold. Once G-CSF treatment had concluded, circulating pmel-1 T cell levels declined slowly across the 14 surviving mice to an average of 3.6 x 10^4^ cells/mL (range = 2.0 x 10^2^ to 1.3 x 10^5^ cells/mL, a 663.2-fold difference) by days 66 to 70 (Fig. 3a, Fig. S2, l and m). To explain the wide range of pmel-1 T cell levels seen during the post-CR persistence phase, we evaluated correlations between various donor T cell factors (peak expansion level, memory phenotype), host cell factors (T cell, neutrophil and B cell levels) and tumour-related factors (tumour size before and during treatment). Of all these parameters, we found a positive trend between day 2 tumour size and pmel-1 persistence on days 66 to 70, which reached statistical significance in one study (P = 0.0182, n = 7 mice; Study 2) (Fig. S2n).

T cell memory phenotypes were assessed over time by flow cytometry. In mice treated with G2R^+^ T cells + G-CSF, the pmel-1 T cells adopted a predominantly Tem phenotype during the period of G-CSF treatment. A similar result was seen in the mock T cells + IL-2 group during the period of IL-2 treatment. For example, in the experiment shown in Fig. 3b, 94.3% and 85.2% of pmel-1 T cells showed a Tem phenotype on day 4 in the G2R^+^ T cells + G-CSF and mock T cells + IL-2-treated groups, respectively. By day 45 (27 or 25 days after the last G-CSF or IL-2 treatment, respectively), pmel-1 T cells from these two groups exhibited a predominantly T central memory (Tcm; CD44⁺CD62L⁺) phenotype (76.3% and 73.0% of pmel-1 T cells, respectively). By contrast, in vehicle-treated mice that received mock or G2R^+^ T cells, a significantly lower proportion of pmel-1 T cells with Tem phenotype was observed on day 4 (65.2% and 67.3%, respectively), and these mice did not survive to day 45 for further evaluation.

### G-CSF induces reversible myeloid expansion with minimal effects on other circulating host cell types

As expected, G-CSF-treated animals showed a rapid and reversible increase in circulating neutrophils during the period of G-CSF administration. For example, in the experiment shown in Fig. 3c, neutrophil levels were 17.0-fold higher on day 16 in mice treated with G2R^+^ T cells + G-CSF compared to the G2R^+^ T cells + vehicle group, consistent with prior studies of G-CSF-treated mice^50,51^. After G-CSF treatment ended, neutrophil counts returned to baseline levels. In vehicle-treated groups, neutrophils increased modestly (2.9-fold) from day 2 to day 16, at which point most animals were euthanized due to tumour burden.

Beyond neutrophils, G-CSF had only modest, reversible effects on host immune cells. As reported by others^52^ and observed in the representative experiment shown in Fig. 3a, circulating NK cells in the G-CSF-treated groups were modestly elevated on day 16 (2.3-fold relative to vehicle-treated groups) (Fig. S3a). Likewise, host CD4^+^ and CD8^+^ T cell counts were approximately 1.3-fold higher in the G-CSF-versus vehicle-treated groups (Fig. 3, d and e). Both NK and T cell counts returned to baseline once G-CSF treatment ended. CD62L expression on host CD8^+^ T cells decreased transiently during G-CSF treatment (Fig. 3f), another previously described effect^53^. In contrast to the predominantly Tcm phenotype of G2R+ pmel-1 T cells at day 80 (Fig. 3b), host CD8^+^ T cells had a predominantly naive phenotype (82.5%) with a minor Tcm subpopulation (12.5%). In all treatment groups, B cell counts were largely unchanged over the course of the study (Fig. S3b).

None of the treatment groups showed any overt toxicities, including skin or coat de-pigmentation (vitiligo), alopecia, or overt vision impairment (indicative of ocular autoimmunity) which have been reported in other immunotherapy studies involving the B16-F10 model^54–56^. Body weights remained within a healthy range in all cases (Fig. S3c).

To assess the effects of treatment on systemic cytokine levels, plasma samples collected on day 4 post-ACT were subjected to 36-plex cytokine array analysis. The mean concentrations of four cytokines differed significantly across treatment groups: CXCL9 (P < 0.0001), CXCL10 (P = 0.0171), IL-6 (P = 0.0067) and IL-13 (P = 0.0204) by Welch’s ANOVA (Fig. 4a). Post hoc comparisons revealed that mice receiving G2R^+^ pmel-1 T cells + G-CSF had significantly elevated levels of CXCL9 compared to most other groups (Fig. S3d), consistent with the ability of G2R to induce CXCL9 production by T cells (Fig. 1f). CXCL10, IL-6 and IL-13 trended higher in mice treated with G2R^+^ T cells + G-CSF or IL-2, but these differences did not reach statistical significance (Fig. S3, e to g).

**Figure 4.**
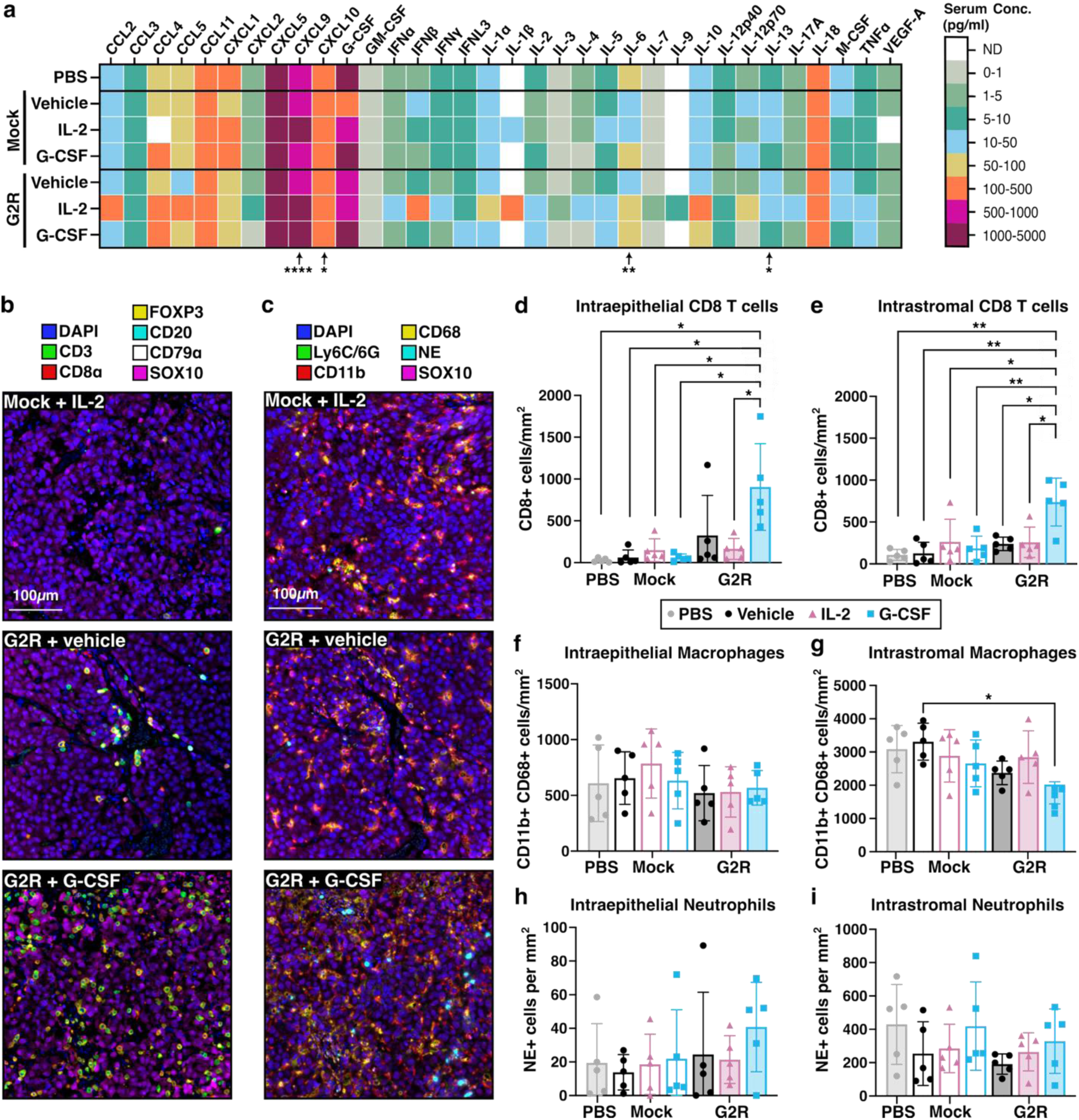
Treatment with G2R^+^ T cells + G-CSF induces systemic CXCL9 and markedly increased tumour infiltration by CD8^+^ T cells. Plasma and tumours (n = 5 per group) were collected on day 4 post-ACT from B16-F10 tumour-bearing mice from the indicated treatment groups. **(a)** Heat map showing the mean levels of the indicated cytokines in plasma for the indicated treatment groups. Not shown are cytokines that were below the limit of detection (IL-15 and LIF). Welch’s ANOVA tests were performed for each cytokine, and significant differences are indicated by asterisks: CXCL9 P < 0.0001; CXCL10 P = 0.0171; IL-6 P = 0.0067; and IL-13 P = 0.0204. **(b-i)** Tumours were analysed by multiplex immunofluorescence to detect: **(b)** cell nuclei (DAPI), CD3, CD8, FOXP3, CD20, CD79a, and SOX10; and **(c)** cell nuclei (DAPI), LY6C/6G, CD11b, CD68, neutrophil elastase (NE), and SOX10. Representative images are shown. **(d-i)** The indicated cell types were quantified in the epithelial and stromal tumour compartments: **(d, e)** CD8^+^ T cells; **(f, g)** macrophages; **(h, i)** neutrophils. Panels **d-i**: unpaired t test with Welch’s correction. *P < 0.05; **P < 0.01; ***P < 0.001; ****P < 0.0001. Data are presented as mean +/-SD of 5 biological replicates. Animals were treated with PEG-GCSF or vehicle on days 0 and 2, or IL-2 daily on days 0 to 3.

### Treatment with G2R^+^ T cells + G-CSF enhances CD8^+^ T cell infiltration into tumours

To assess the impact of different treatments on the tumour microenvironment, we applied multiplex immunofluorescence (mpIF) to tumours harvested on day 4 post-ACT (Fig. 4, b and c), a timepoint when pmel-1 T cell expansion was highest in peripheral blood yet tumour regression was incomplete. Compared to control mice that received PBS alone, tumours from mice treated with mock-transduced T cells and either vehicle or G-CSF exhibited a modestly higher (mean = 2.2-fold) density of intraepithelial CD8^+^ T cells (Fig. 4d). An even higher increase (mean = 11.6-fold) was seen after treatment with G2R^+^ T cells + vehicle. More strikingly, treatment with G2R^+^ T cells + G-CSF induced a 32.3-fold increase in intraepithelial CD8^+^ T cells, which coincided with high levels of circulating pmel-1 T cells (mean = 5.1 x 10^5^ cells/mL) in matched blood samples (Fig. S4a). Intrastromal CD8^+^ T cell density followed a similar pattern, with a 6.9-fold increase seen in the G2R^+^ T cells + G-CSF treatment group relative to control mice that received PBS alone (Fig. 4e).

In contrast to the increased CD8^+^ T cell densities seen by mpIF, there were no significant changes in the densities of intraepithelial or intrastromal CD3^+^CD8^-^T cells (presumptive CD4^+^ T cells), CD8^-^FoxP3^+^ T cells (presumptive Tregs), CD20^+^CD79a^+^ B cells, or CD20^-^CD79a^+^ plasma cells across any of the treatment groups (Fig. S4, b to e). Furthermore, we observed no significant differences in the densities of intraepithelial or intrastromal CD68^+^CD11b^+^ macrophages or neutrophil elastase-expressing neutrophils in the G2R^+^ T cells + G-CSF group compared to the mock-transduced + vehicle group (Fig. 4, f to i), despite the marked increases in circulating neutrophils seen in the former group (Fig. S4f).

### G2R^+^ T cells are associated with a long-term protective memory population that can be boosted by G-CSF re-treatment

Of the 14 mice across three experiments that achieved CR after treatment with G2R^+^ T cells + G-CSF, 10 (71.4%) rejected a second challenge of B16-F10 cells delivered 74 to 98 days after the initial (and only) infusion of G2R^+^ T cells (Fig. 5a, Table S2). As exemplified by animal #1 in Fig. 5b, these 10 mice had a 9.6-fold higher average level of circulating pmel-1 T cells prior to tumour re-challenge compared to the remaining four mice that failed to reject tumour re-challenge (Fig. 5c). The proportion of pmel-1 T cells with a Tcm phenotype also trended higher in mice that rejected tumour re-challenge compared to those that did not (84.4% versus 62.6%, respectively) (Fig. 5d).

**Figure 5.**
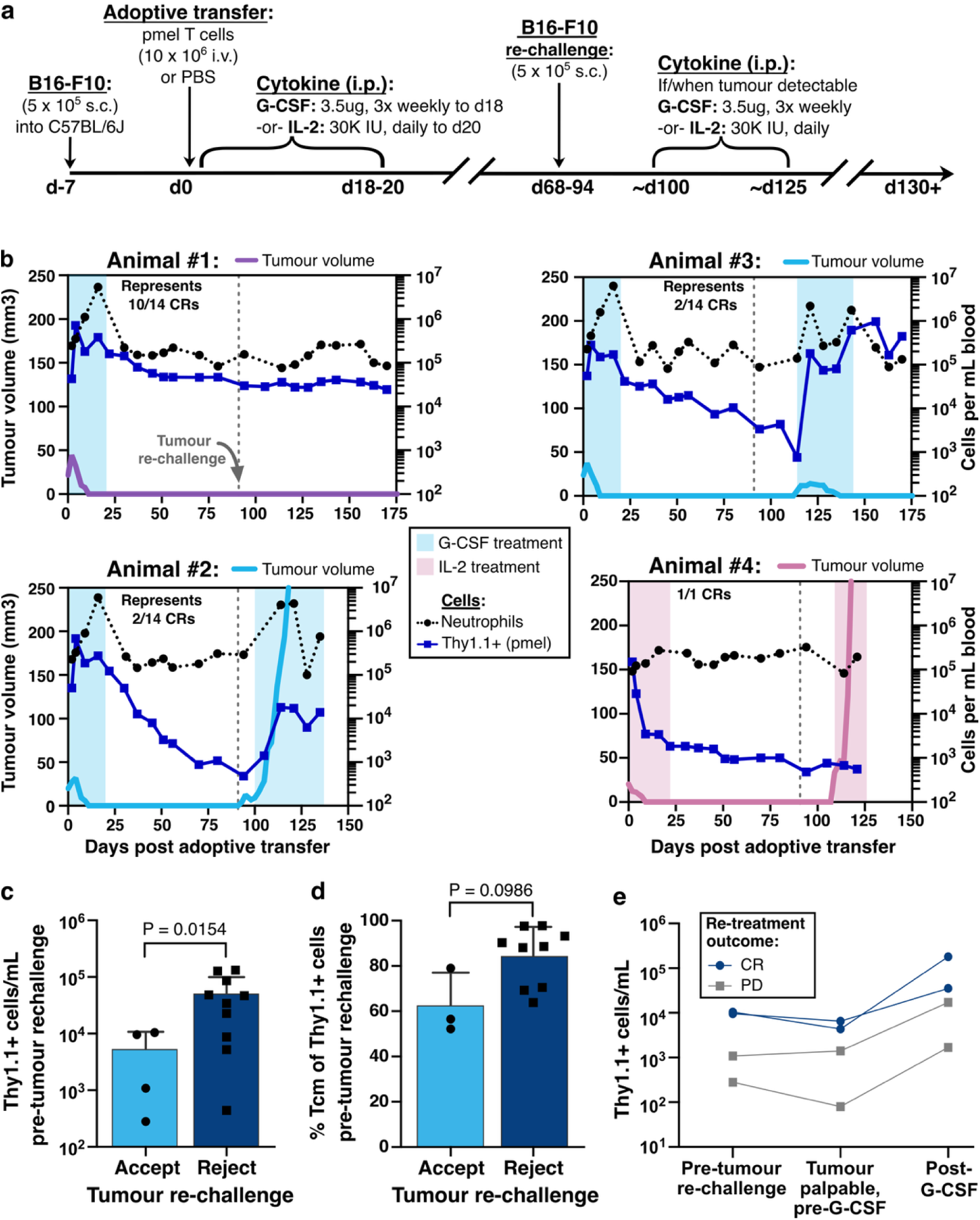
G2R-expressing T cells confer strong resistance to tumour re-challenge and when necessary can be re-expanded with G-CSF administration. **(a)** Experimental schema for the tumour re-challenge experiments involving mice that exhibited CR after initial treatment with G2R^+^ pmel-1 T cells + G-CSF (n = 14) or mock-transduced pmel-1 T cells + IL-2 (n = 1). Mice were re-challenged with a subcutaneous injection of 5 x 10^5^ B16-F10 cells on day 74 to 98 (depending on the experiment) after the first (and only) ACT. If tumours engrafted, mice were re-treated with G-CSF or IL-2 (as per their original treatment group) until CR or humane endpoint. **(b)** Summary of results for four representative mice, showing tumour volume, circulating pmel-1 T cells, neutrophils, and re-treatment interval with G-CSF (Animals #2 and #3) or IL-2 (Animal #4). Animal #1 rejected tumour re-challenge and therefore did not require G-CSF re-treatment. **(c)** Number and **(d)** Tcm phenotype of circulating pmel-1 T cells pre-tumour re-challenge, stratified by mice that rejected vs. accepted the re-challenge. Memory phenotype could only be determined for 12/14 mice with sufficient pmel-1 counts. **(e)** Number of circulating pmel-1 T cells in four mice that were re-treated with G-CSF. Three timepoints are shown: pre-tumour re-challenge; when tumour was palpable but before re-treatment with G-CSF; and after at least 15 days of G-CSF re-treatment. Colours distinguish mice that exhibited CR (blue lines) versus progressive disease (PD, gray lines) following G-CSF re-treatment. Panels **c-d**: unpaired t testwith Welch’s correction.

The four mice that failed to reject tumour re-challenge were re-treated with G-CSF on the first indication of tumour growth, and G-CSF was continued until CR was observed or tumours reached the prescribed endpoint. Upon G-CSF re-treatment, all four mice showed an increase in circulating pmel-1 T cells (Fig. 5e). Of these four mice, the two that achieved the lowest pmel-1 T cell levels after G-CSF re-treatment experienced progressive tumour growth (e.g., animal #2 in Fig. 5b). In contrast, the two that achieved the highest pmel-1 T cell levels after G-CSF re-treatment exhibited second, durable CRs within 6 to 21 days (e.g., animal #3 in Fig. 5b). As a comparator to G-CSF re-treatment, the one mouse that had originally achieved a CR after treatment with mock-transduced T cells + IL-2 (Fig. 2b) failed to reject a second tumour challenge (animal #4, Fig. 5b). This animal received a second round of IL-2 treatment; however, the pmel-1 T cells failed to expand or induce tumour regression (Fig. 5b and Table S2). Thus, in addition to providing protective memory responses in the majority (10/14, 71.4%) of mice, G2R engineering enables on-demand, G-CSF-induced re-expansion of tumour-specific T cells without need for additional ACT or any other form of therapy.

## Discussion

TIL therapy for metastatic melanoma is transitioning from an investigational to standard-of-care treatment option, yet widespread application is held back by the suboptimal efficacy of current regimens and the often-serious toxicities associated with lymphodepletion and high-dose IL-2^8,9^. Here, we show in the B16-F10 murine model of melanoma that the need for lymphodepletion and high-dose IL-2 can be eliminated by engineering T cells to express a single gene product, G2R, which generates an IL-2-like signal in response to G-CSF, an approved, well-tolerated cytokine. This strategy enables profound increases in T cell expansion and persistence, leading to durable, complete regression of established melanoma in over 75% of mice. Moreover, G2R-expressing T cells provide long-term protective immunity against tumour re-challenge, and when needed, can be re-expanded to therapeutically effective levels with repeat G-CSF treatment alone. To our knowledge, no other immunotherapy achieves an equivalent level of efficacy in this widely used model without the need for lymphodepletion, exogenous IL-2 or any other co-therapy. Furthermore, the approach has high feasibility for clinical translation.

As discussed earlier, a wide variety of strategies have been investigated to improve the safety and efficacy of TIL therapy by reducing the need for lymphodepletion and/or high-dose IL-2. These include designer IL-2/IL-15 superagonists^12–14^; cis-acting IL-2/IL-21 fusion proteins that selectively engage T cells co-expressing both receptors^16^; cytokine switch receptors^17–19^; cytokine-armoured T cells^20–22^; orthogonal IL-2/IL-2Rβ pairs^24^; and acetazolamide-regulated expression of membrane-bound IL-15^30^. While these approaches have merit, they generally require the production and clinical validation of a novel biologic and/or complex cell engineering strategy. Furthermore, several of these approaches bring the risk of immunogenicity of the engineered cytokine and/or receptor, and several have documented toxicities^16,23,26^. In contrast, with G2R technology, G-CSF is used in an unmodified clinical formulation and pairs with a single gene product containing a single modification (i.e., the fusion site between the JAK binding domain of G-CSFR and the C-terminal region of IL-2Rβ). This reduces manufacturing complexity, risk of immunogenicity, cost of goods and regulatory burden. Moreover, G2R can be activated on-demand through the provision of exogenous G-CSF, thereby conferring clinician control.

G2R technology also has a favourable safety profile in mice. As expected, G-CSF induced transient neutrophilia, which fully resolved after G-CSF cessation, without observable weight loss or any other signs of systemic toxicity. Additionally, there were no significant increases in intratumoural neutrophils or macrophages, despite the systemic myeloid response. Plasma cytokine profiles were also favourable. Consistent with the CXCL9 secretion we observed from G2R^+^ T cells in vitro, mice treated with G2R^+^ T cells + G-CSF showed an increase in circulating CXCL9. IL-6 showed a modest but insignificant increase in the G2R^+^ T cells + G-CSF group (reaching an average of 56.7 pg/mL), consistent with the expected systemic footprint of an actively proliferating and cytolytic T cell population; however, this is an order of magnitude below the median peak IL-6 levels in patients experiencing cytokine release syndrome after CAR-T cell therapy (606 pg/mL)^57^. Notably, we did not see upregulation of sIL-2Rα or IL-5, which are associated with high-dose IL-2 treatment^58,59^ and some engineered cell therapy platforms^26^. Finally, we did not observe any of the toxicities reported in other immunotherapy studies involving the B16-F10 model, including skin or coat de-pigmentation (vitiligo), alopecia, or overt vision impairment (indicative of ocular autoimmunity)^54–56^. Thus, G2R technology can promote potent anti-tumour responses without causing apparent damage to healthy tissues expressing the gp100 target antigen.

Despite these advantages, the use of G-CSF also brings special considerations. The first concerns baseline levels of G-CSF, which could theoretically promote uncontrolled T cell proliferation, possibly at the expense of normal myelopoiesis. Serum G-CSF levels are in the range of 15-22 pg/mL in healthy humans^36^, 14-300 pg/mL in cancer patients^60–62^, and up to 4,300 pg/mL in certain inflammatory conditions^61^. Thus, apart from extreme inflammatory conditions, G-CSF levels are one to two orders of magnitude below that required to stimulate G2R-induced T cell proliferation in vitro. Accordingly, we saw no evidence of expansion of G2R^+^ pmel-1 T cells in mice in the absence of exogenous G-CSF. Several types of human tumours produce G-CSF^63–65^, which could lead to higher levels in the tumour microenvironment; however, this could in fact be a favourable factor, as it would promote T cell proliferation and anti-tumour activity locally, where it is most needed. Another consideration is that G-CSF is the mainstay pharmaceutical agent used to treat neutropenia in cancer patients, including during TIL therapy^9,31^, and re-purposing it to also stimulate TIL results in a more complex pharmacology and clinical application. Importantly, we showed in the B16-F10 model that G-CSF can simultaneously enhance both neutrophil and G2R^+^ T cell counts in a safe and reversible manner. In a scenario where more neutrophils were needed but stimulation of TIL was contraindicated, granulocyte-macrophage colony-stimulating factor (GM-CSF) could potentially be used to selectively stimulate myelopoiesis without impacting G2R^+^ T cells. Finally, G2R technology offers the potential to reduce or eliminate the need for lymphodepletion, which may in turn reduce the incidence of neutropenia and the need to use G-CSF for that purpose.

Several factors could potentially contribute to the extraordinary T cell expansion and anti-tumour efficacy seen with G2R technology. First, G2R signalling may generate a more potent intracellular signal compared to the native IL-2R owing to the unique engagement of JAK2, STAT4 and/or potentially other unknown signalling intermediaries. Notably, however, G2R and the wildtype IL-2R and IL-15R induced similar levels of in vitro T cell proliferation and cytokine production (including IFN-γ, which is downstream of STAT4^66^), suggesting that biochemical signalling differences may not be responsible for the observed in vivo differences. Second, the stimulatory effect of G-CSF on host myeloid and/or NK cells could work synergistically with G2R signalling to enhance anti-tumour immunity. This explanation, however, was not supported by the experiment in which mice were treated with mock-transduced T cells + IL-2 + G-CSF, which induced myeloid and NK cell expansion yet without an enhanced anti-tumour effect. Third, unlike IL-2 administration^67,68^, G2R + G-CSF signalling should not induce a Treg response, thereby allowing unimpeded expansion and anti-tumour activity of pmel-1 T cells. While this is a compelling hypothesis, we note that IL-2 treatment failed to induce increased FoxP3^+^ T cell densities in the tumour microenvironment in our experiments. Fourth, the remarkable expansion of G2R^+^ pmel-1 T cells may be largely attributable to the exclusive delivery of an IL-2 signal to the pmel-1 T cell population without concurrent stimulation of host T cells and the competition for space and resources that might result. Fifth, and somewhat opposite to the preceding idea, the lack of lymphodepletion in the G2R^+^ T cells + G-CSF treatment regimen could have enabled host T cells, NK cells and potentially other immune cells to contribute to the anti-tumour response, resulting in greater efficacy^10,11^. Indeed, our multiplex immunofluorescence study revealed substantial infiltration of the tumour microenvironment by host T cells and myeloid cells.

In addition to mediating a potent anti-tumour response, treatment with G2R^+^ T cells + G-CSF led to long-term persistence and protective immunity, which was associated with a shift from a predominantly Tem to Tcm phenotype after G-CSF withdrawal and tumour clearance. This contrasts with traditional IL-2-supported ACT, where IL-2 can drive terminally differentiated or exhausted T cell phenotypes, which can hinder persistence and efficacy^69^. However, for reasons which remain unclear, the level of persistent pmel-1 T cells varied by over 3,100-fold on days 16-17 and 660-fold on days 66-70 in mice that had exhibited CR. In exploratory analyses, higher peak pmel-1 T cell numbers were positively associated with a proportionately higher Tem phenotype, a trend that was significant in one of three studies. Additionally, there was a trend toward larger tumour volume on day two post-ACT being associated with greater pmel-1 T cell persistence on day 80, which also reached statistical significance in one study. One possible explanation is that greater tumour burden provides increased antigen stimulation, which, when paired with G-CSF-driven signalling through G2R, may increase the size of the memory T cell pool that is formed. Whether tumour burden is a direct determinant of G2R^+^ T cell persistence will need to be tested directly in future studies.

Irrespective of the underlying cause, the level of persisting pmel-1 T cells was significantly associated with the ability to reject a second tumour challenge. Thus, another future direction is to determine whether prophylactic administration of G-CSF during the remission phase can increase T cells to a “protective setpoint” in all mice. An alternative option, as shown here, is to re-treat with G-CSF if and when needed to treat second tumours (or recurrent tumours in a clinical scenario). Yet even this option appeared to be more successful in mice that had higher pmel-1 T cell levels prior to tumour re-challenge, suggesting that the optimal approach might be a combination of G-CSF prophylaxis and as-needed re-treatment. In a clinical setting, such an approach could be done on a personalized basis simply by tracking the level of G2R^+^ T cells by flow cytometry, as shown in our human T cell experiments.

Despite the strong anti-tumour efficacy reported here, a limitation of this study is the use of a single, relatively immunogenic mouse tumour model with a homogeneously expressed target antigen (gp100). Moreover, ACT was performed with large numbers of monoclonal, tumour-specific CD8^+^ T cells which do not reflect the heterogeneity of human TIL with respect to CD4:CD8 content, TCR diversity, tumour specificity, or states of activation, differentiation and dysfunction. That said, the B16-F10/pmel-1 model is among the most widely used for the development and validation of T-cell based immunotherapies^70–74^ and therefore provides important benchmarks for the exceptional efficacy and safety of our approach. Nonetheless, a future direction is to investigate G2R technology in additional tumour and T cell models to establish broader applicability. Indeed, in addition to initiating the clinical translation of G2R-engineered TIL therapy for advanced melanoma and other solid tumours, we are investigating the use of G2R in CAR-T cell therapy, where cytokine signalling also plays a critical role in amplifying T-cell expansion, activity and persistence^38,75,76^.

## Methods

### Cell lines

The HEK293T cell line (Cat. CRL-3216) and B16-F10 mouse melanoma cell (Cat. CRL-6475) line were purchased from American Type Culture Collection (ATCC, Manassas, Virginia,) and maintained according to the manufacturer’s recommendations. The Platinum-Eco RV cell line (RV-101) was purchased from Cell Biolabs (San Diego, California) and maintained according to manufacturer’s recommendations.

### Cytokine preparation and administration

IL-2 (Proleukin; Iovance, San Carlos, California, DIN 02130181, lot. M002595) was procured from institutional pharmacy (BC Cancer Victoria, British Columbia). Drug was resuspended in PBS to 300,000 IU/mL and frozen at -80°C until use. Once thawed, aliquots were stored at 4°C for a maximum of three days during use. 30,000 IU (100 µL) was administered by intraperitoneal (IP) injection once daily for a total of 20 days unless otherwise noted.

PEG-GCSF (Neulasta, Amgen, Thousand Oaks, California, Cat. 1-01190-01, 6mg, lot 1161444) was procured from institutional pharmacy (BC Cancer Victoria, British Columbia) and diluted to a final concentration of 35 µg/mL in vehicle (10 mM Sodium Acetate, 5% (w/v Sorbitol), 0.004% Polysorbate-20, pH 4.0). 3.5 µg (100 µL) was administered by IP injection three times per week for a total of 18 days unless otherwise noted. 100 µL of vehicle was administered by IP injection three times per week for a total of 18 days unless otherwise noted.

All mice were closely monitored for 10 minutes after each injection for signs of distress and side effects.

### Lentiviral vector production

Lentiviral vectors were produced by transient transfection of HEK293T cells (ATCC, Manassas, Virginia, Cat. CRL-3216) using Lipofectamine 3000 (Thermo Fisher, Waltham, Massachusetts, Cat. L3000015), according to the manufacturer’s instructions. Briefly, HEK293T cells were maintained in Opti-MEM reduced serum medium with Glutamax supplement (Gibco, Brooklyn, New York, Cat. 51985034) + 5% fetal bovine serum (FBS; Sigma-Aldrich, St. Louis, Missouri, Cat. F1051) + 0.2mM sodium pyruvate (Cytiva Life Sciences, Marlborough, Massachusetts, Cat. SH30239.01) under standard cell culture conditions and transfected with three plasmids: the lentiviral transfer vector encoding the gene of interest, a packaging plasmid encoding GAG/POL and REV, and a plasmid encoding the vesicular stomatitis virus glycoprotein (VSV-G) envelope protein. Plasmid DNA and Lipofectamine 3000 complexes were prepared using the accompanying P3000 reagent and added to the cells. Following transfection, the culture medium was replaced. Lentivirus-containing culture supernatants were collected at 24 and 48 hours, clarified to remove cellular debris, and filtered through a 0.45µm filter. Viral preparations were concentrated, aliquoted, and stored at -80°C until use.

### Human T cell isolation, activation, lentiviral transduction, expansion and flow cytometry

Human T cells were derived from healthy donor leukopacks (Stemcell Technologies, Vancouver, British Columbia, Cat. 70500.2). All leukopacks were commercially sourced with informed consent being obtained by the supplier. CD3^+^ T cells were isolated using a CD3^+^ T cell negative selection isolation kit (Stemcell Technologies, Vancouver, British Columbia, Cat. 17951) and cryogenically preserved in a 50% solution of Cryostore (Stemcell Technologies, Vancouver, British Columbia, Cat. 07930). Cells were then thawed and cultured in TexMACS human T cell medium (Miltenyi Biotec, Bergisch Gladbach, Germany, Cat. 130-097-196) + 3% Human Serum (Sigma-Aldrich, St. Louis, Missouri, Cat. H4522) at 1 x 10^6^ per well and stimulated with TransAct (Miltenyi Biotec, Bergisch Gladbach, Germany, Cat. 130-111-160) and human IL-2 (300 IU/mL). The next day, cells were transduced as indicated using lentivirus encoding G2R at a multiplicity of infection (MOI) of 5. Expansion was continued using either IL-2 (300 IU/mL) or switching to G-CSF (100 ng/mL) (Apotex, Toronto, Ontario, DIN 02454548) from day 2 onwards. Cells were expanded a total of 13 days for most downstream assays, or up to 20 days for BrdU analysis. Mock-transduced T cells were cultured using the same procedures as G2R-transduced T cells but excluding the lentiviral vector. Fluorescently labelled antibodies were used to quantify surface expression of proteins of interest. All antibodies used were titrated to determine optimal concentration for detection as detailed in Table S3. Antibodies were diluted using cell staining buffer (BioLegend, San Diego, California, Cat. 420201) in addition to Brilliant Stain Buffer Plus (BD Biosciences, Franklin Lakes, New Jersey, Cat. 563794) and incubated with human CD3^+^ T cells for 30 minutes at 4°C in the dark. Cells were washed twice using PBS + 2% FBS (Sigma-Aldrich, St. Louis, Missouri, Cat. F1051) prior to resuspension in cell staining buffer. Samples were acquired on a Cytek Aurora 3-laser cytometer using SpectroFlo software (Cytek Biosciences, Fremont, California). Dead cells were excluded from analysis using Zombie NIR viability dye (BioLegend, San Diego, California, Cat. 423106). Analysis performed using SpectroFlo software (Cytek Biosciences, Fremont, California).

### BrdU proliferation assay of human T cells

BrdU incorporation was assessed using the BD Pharmingen^TM^ FITC BrdU Flow Kit (BD Biosciences, Franklin Lakes, New Jersey, Cat. 559619) according to the manufacturer’s instructions. Human CD3⁺ T cells were expanded as described above and harvested on day 13 or day 20 of expansion. Cells were washed and resuspended in TexMACS human T cell medium (Miltenyi Biotec, Bergisch Gladbach, Germany, Cat. 130-097-196) + 3% Human Serum (Sigma-Aldrich, St. Louis, Missouri, Cat. H4522) at 2 × 10⁶ cells/mL, then plated at 1 × 10⁶ cells per well in 48-well plates. Cells were cultured in the presence of recombinant human IL-2 (300 IU/mL), G-CSF (100 ng/mL), recombinant human IL-15 (10 ng/mL) (Miltenyi Biotec, Bergisch Gladbach, Germany, Cat. 130-095-760), or no added cytokine and incubated for 48 h at 37°C with 5% CO₂. BrdU was added 32 hours after plating at the concentration recommended by the manufacturer, and cells were incubated for the remaining 16 hours. For the dose-response assay, G-CSF concentrations were in half-log increments from 0.001 ng/mL to 1000 ng/mL.

At the end of the incubation period, cells were harvested and stained with antibodies against cell surface markers (CD3, CD4, CD8, and G-CSFR), followed by viability staining with Zombie NIR (BioLegend, San Diego, California, Cat. 423106). Cells were subsequently fixed, permeabilized, treated with DNase, and stained with anti-BrdU-FITC in accordance with the kit protocol. DNA content was assessed by 7-AAD staining. Samples were acquired on a Cytek Aurora 3-laser cytometer using SpectroFlo software (Cytek Biosciences, Fremont, California). Dead cells were excluded from analysis using Zombie NIR viability dye (BioLegend, San Diego, California, Cat. 423106). Analysis performed using SpectroFlo software (Cytek Biosciences, Fremont, California).

### Human T cell stimulation and western blotting to assess intracellular signalling

Human T cells were washed three times in PBS, prior to resting overnight in cytokine-free complete medium (TexMACS medium (Miltenyi Biotec, Bergisch Gladbach, Germany, Cat. 130-097-196) + 3% Human Serum (Sigma-Aldrich, St. Louis, Missouri Cat. H4522) before cytokine stimulation. Cells were adjusted to 2 x 10^6^ cells/ml and stimulated with either IL-2 (300 IU/mL), IL-15 (10 ng/mL), or G-CSF (100 ng/mL) for 20 minutes. Unstimulated cells were used as a baseline control. Cells were then rapidly collected by centrifugation at 4°C and washed with ice-cold wash buffer (10 mM HEPES pH 7.9, 1 mM MgCl2, 0.05 mM EGTA pH 8.0, 0.5 mM EDTA pH 8.0, 1 mM DTT) containing protease and phosphatase inhibitors (Thermo Fisher, Waltham, Massachusetts, Cat. A32959). Cell pellets were lysed at 1 x 10^6^ cells/10 μL lysis buffer (wash buffer with 0.2% NP-40). Lysates were incubated on ice for 10 minutes and clarified by centrifugation at 18,000 x g for 10 minutes at 4°C. Cytoplasmic supernatants were collected. The remaining nuclear pellets were resuspended at 1 x 10^6^ cell equivalents/4 μL of lysis buffer (wash buffer with 0.42 M NaCl, 20% glycerol) and incubated on ice for 30 minutes with mixing every five minutes. Lysates were centrifuged at 18,000 x g for 20 minutes at 4°C, and supernatants were collected.

Protein samples were diluted in NuPAGE reducing agent (Thermo Fisher, Waltham, Massachusetts, Cat. NP0009) and LDS sample buffer (Thermo Fisher, Waltham, Massachusetts, Cat. NP0007) and denatured at 70°C for 10 minutes. Equivalent amounts of total protein (1.67 x 10^6^ cell equivalents of nuclear lysate per lane, 1.11 x 10^6^ cell equivalents of cytoplasmic lysates) were resolved by SDS-PAGE using 4-12% polyacrylamide gels (Thermo Fisher, Waltham, Massachusetts, Cat. NP0321BOX) at 200V for 50 minutes. Proteins were transferred to 0.2 μm nitrocellulose membranes using semi-dry transfer at 25V 2.5A for 12 minutes with an Invitrogen Powerblotter. Successful transfer was assessed by Ponceau S staining. Membranes were washed and blocked in LI-COR Intercept TBS Blocking Buffer (LI-COR Bio, Lincoln Nebraska, Cat. NC1660550) for 1 hour at room temperature.

Membranes were incubated overnight at 4°C with primary antibodies directed against various target proteins: Phospho-JAK1 (Cell Signaling Technology, Danvers, Massachusetts [CST], 74129S), Phospho-JAK2 (CST 8082S), Phospho-JAK3 (CST 5031S), Phospho-P70S6K (CST 9204S), Phospho-S6 (CST 2211S), Phospho-AKT (CST 9275S), Phospho-Shc (CST 2434S), Phospho-ERK (CST 9101S), Phospho-STAT1 (CST 9167S), Phospho-STAT3 (CST 9138S), Phospho-STAT4 (CST 5267S), Phospho-STAT5 (CST 9359S), Beta-Actin (CST 3700S), and Histone H3 (CST 3638S), all at 1:1,000 dilution. Following three washes in Tris-buffered saline containing 0.05% Tween20 (TBS-T) (Thermo Fisher, Waltham, Massachusetts, Cat. 28360), membranes were incubated with fluorescently labelled secondary antibodies; IRDye 800CW goat anti-rabbit (LI-COR Bio, Lincoln Nebraska, NC9401842), or IRDye 680RD goat anti-mouse (LI-COR Bio, Lincoln Nebraska, NC0252290) at 1:10,000 dilution in TBS-T for 60 minutes at room temperature. Membranes were then washed three times with TBS-T, followed by one wash in TBS. Fluorescent signals were detected using a LI-COR Odyssey M imaging system and LI-COR Acquisition 2.1 software (LI-COR Bio, Lincoln Nebraska).

### Mice

All animal studies were conducted according to protocols approved by the University of Victoria’s Animal Care Committee. Six- to eight-week-old female C57BL/6J mice (Strain #000664, RRID: IMSR_JAX:000664), and B6.Cg-*Thy1^a^*/Cy Tg(TcraTcrb)8Rest/J (pmel-1) mice (Strain #005023, RRID: IMSR_JAX:005023) were purchased from Jackson Laboratories (Bar Harbor, Maine). Mice were housed at the University of Victoria’s Animal Care Unit in accordance with institutional guidelines and practices. Animals were maintained under standard pathogen-free conditions at 22°C with a 12 h light/dark cycle. Experiments were carried out using eight- to twelve-week-old animals.

### Tumour engraftment and randomization

B16-F10 tumour cells (5 x 10^5^) were injected subcutaneously into the right flank of recipient C57BL/6J mice. Tumours were measured three times per week using calipers. Investigators were blinded to the treatment groups during measurements. Tumour volume was estimated using the following formula: (length x width^2^) x 0.5 = mm^3^. Tumours were left to grow for seven days before ACT. On Day 7 post tumour implant, mice were randomized into treatment groups based on average tumour volume. Mice were euthanized when tumour volume reached 1,000 mm³ or earlier if humane endpoint criteria were met, including a deteriorating body score, weight loss exceeding 20%, and necrotic or infected tumours. Euthanasia was performed by isoflurane inhalation (4-5%) followed by cervical dislocation.

### Mouse pmel-1 T cell isolation, retroviral transduction, culture, flow cytometry and adoptive transfer

Female pmel-1 mice were used as donors for ACT. Spleens were harvested from euthanized mice, and CD8⁺ T cells were isolated by negative selection using the EasySep Mouse CD8⁺ T Cell Isolation Kit (Stemcell Technologies, Vancouver, British Columbia, Cat. 19853), yielding >90% purity. Immediately following isolation, purified CD8⁺ T cells were cultured at 1 × 10⁶ cells/mL in mouse T cell medium (DMEM high glucose (Gibco, Brooklyn, New York, Cat. 11-965-092)) + 10% FBS (Sigma-Aldrich, St. Louis, Missouri, Cat. F1051) + 0.1% β-mercaptoethanol (Sigma-Aldrich, St. Louis, Missouri, Cat. 444203) and activated with soluble anti-CD3 (1 μg/mL) (BD Biosciences, Franklin Lakes, New Jersey, Cat. 553058), anti-CD28 (0.5 μg/mL) (BD Biosciences, Franklin Lakes, New Jersey, Cat. 553294), and recombinant human IL-2 (300 IU/mL). Cells were maintained at 37 °C with 5% CO₂ throughout culture.

The retroviral plasmid, MIGR1 was a gift from Warren Pear (Addgene plasmid # 27490; http://n2t.net/addgene:27490; RRID:Addgene_27490). The G2R cDNA was cloned into the MIGR1 retroviral vector in place of the IRES-GFP fragment. Retroviral particles encoding the murine G2R construct were generated using the Platinum-Eco packaging cell line (Cell Biolabs, San Diego, California). Platinum-Eco cells were maintained under standard selection conditions and transiently transfected using Lipofectamine 3000 (Thermo Fisher, Waltham, Massachusetts, Cat. L3000015) according to the manufacturer’s instructions. Viral supernatants were collected at 24 and 48 hours post-transfection, filtered, supplemented with polybrene (Sigma-Aldrich, St. Louis, Missouri, Cat. TR-1003-G) and IL-2 and used immediately for transduction.

Activated pmel-1 T cells were transduced by spinfection on two consecutive days using freshly collected viral supernatant. Where indicated, mock-transduced T cells underwent identical activation and culture conditions in the absence of viral supernatant. Following transduction, cells were cultured for an additional three days in mouse T cell medium containing IL-2 (300 IU/mL). On day 5 of culture, cells were harvested, washed, and counted using Acridine orange/Propidium iodide stain (Logos Biosystems, Gyeonggi-do, South Korea, Cat. F23001) and Luna-FX7 Automated Cell Counter (Logos Biosystems, Gyeonggi-do, South Korea). Cells were resuspended in PBS, and 1 × 10⁷ viable T cells (100 μL) were administered to recipient mice by intravenous tail vein injection on day 7 following tumour injection.

Fluorescently labelled antibodies were used to quantify surface expression of proteins of interest. All antibodies used were titrated to determine optimal concentration for detection and are detailed in Table S4. Antibodies were diluted using cell staining buffer (BioLegend, San Diego, California, Cat. 420201) in addition to Brilliant Stain Buffer Plus (BD Biosciences, Franklin Lakes, New Jersey, Cat. 563794) and incubated with mouse peripheral blood samples for 30 minutes at 4°C in the dark. Cells were washed twice using PBS + 2% FBS (Sigma-Aldrich, St. Louis, Missouri, Cat. F1051) prior to resuspension in staining buffer. Samples were acquired on a Cytek Aurora 3-laser cytometer using SpectroFlo software (Cytek Biosciences, Fremont, California). Dead cells were excluded from analysis using Zombie NIR viability dye (BioLegend, San Diego, California, Cat. 423106). Analysis performed using SpectroFlo software (Cytek Biosciences, Fremont, California).

### Phospho-STAT5 flow cytometry assay

Human or mouse CD3⁺ T cells were expanded as described above and harvested on day 13 (human) or day six (mouse) of expansion. Cells were washed and resuspended in TexMACS human T cell medium (Miltenyi Biotec, Bergisch Gladbach, Germany) at 1 × 10⁶ cells/mL, then transferred to FACS tubes at 0.5 × 10⁶ cells per tube in cytokine-free media and incubated overnight at 37 °C with 5% CO₂. At the end of the incubation period, cells were stained with antibodies against cell surface markers (CD3, CD4, CD8, and G-CSFR), followed by viability staining with Zombie NIR. Cells were then stimulated with cytokine (IL-2 [300 IU/mL] or G-CSF [100 ng/mL]) in technical replicates for 5 to 60 minutes at 37°C, followed by immediate fixation, permeabilization and intracellular staining with antibodies specific for phospho-STAT5 (BD Biosciences, Franklin Lakes, New Jersey, Cat. 562077) or isotype control (BD Biosciences, Franklin Lakes, New Jersey, Cat. 551436). Samples were acquired on a Cytek Aurora 3-laser cytometer using SpectroFlo software (Cytek Biosciences, Fremont, California). Dead cells were excluded from analysis using Zombie NIR viability dye (BioLegend, San Diego, California, Cat. 423106). Analysis performed using SpectroFlo software (Cytek Biosciences, Fremont, California).

### Multiplex cytokine analysis

Human T cells expressing G2R or mock-transduced T cells were collected on day 12 of expansion and washed three times in PBS. Cells were then plated in media containing either IL-2 (300 IU/mL), G-CSF (100 ng/mL), or no cytokine. On day 16, supernatants were harvested from the cultures and shipped to Eve Technologies (Calgary, Alberta) for analysis using the Human Cytokine/Chemokine Panel A 48-Plex Discovery Assay® Array (Cat. HD48A).

For murine cytokine studies, blood was collected into EDTA tubes from a terminal cardiac puncture on day 4 post ACT. Blood was fractionated by centrifugation at 1000 x g for 10 minutes, and plasma layer collected. Plasma was analysed by Eve Technologies (Calgary, Alberta) using the Mouse Cytokine/Chemokine 36-Plex Discovery Assay® Array (Cat. MD36).

### Quantification of tumour-infiltrating immune cells by multiplex immunofluorescence

B16-F10 tumours were collected from mice immediately following euthanasia and stored in 10% formalin solution (Sigma-Aldrich, St. Louis, Missouri, Cat. HT501128-4L) for a minimum of 24 hours prior to further processing. Tumour samples were processed in a TP1020 Carousel tissue processor (Leica, Wetzlar, Germany) using a series of reagent alcohols and xylene (Fisher Scientific, Pittsburgh, Pennsylvania). Tissue was then used to construct tissue microarrays with a manual tissue microarrayer (Beecher Instruments, Sun Prairie, Wisconsin) using triplicate cores per sample. Tissue microarray blocks were sectioned at 4 µm (Microm HM355 microtome) onto glass slides. Slides were deparaffinized using xylene and a dilution series of reagent alcohols, then incubated for 20 minutes in 10% neutral buffered formalin (NBF; MilliporeSigma Canada Ltd., Oakville, ON, Canada), followed by two rinses in distilled water. Antigen retrieval was performed using Diva Decloaker solution in a Decloaking Chamber (Biocare, Concord, California).

Slides were loaded into an Intellipath FLX autostainer (Biocare, Concord, California) for mpIF staining using two antibody panels focused on lymphocyte or myeloid markers. For both panels, endogenous peroxidase activity and non-specific binding were first blocked with Peroxidase-1 (five minutes at room temperature) and Background Sniper (10 minutes at room temperature; Biocare, Concord, California). Staining then proceeded through six sequential rounds, each comprising primary antibody incubation, an HRP-polymer secondary, and an Opal fluorophore-TSA step (Akoya Biosciences, Marlborough, Massachusetts), followed by microwave treatment in AR6 or AR9 antigen retrieval solution (Akoya Biosciences, Marlborough, Massachusetts) to strip the antibody complex while preserving the covalently bound fluorophore.

For the lymphocyte panel (Fig. 4b), primary antibodies were applied in the order CD3 (clone SP7, Abcam, Cambridge, England; Da Vinci Green Diluent, 30 minutes, room temperature), FoxP3 (clone FJK-16s, Fisher Scientific, Pittsburgh, Pennsylvania; Rat 2-Step HRP, 5 minutes probe/10 minutes polymer; Biocare, Concord, California), CD20 (clone E3N7O, Cell Signaling Technology, Danvers, Massachusetts), CD8α (clone D4W2Z, Cell Signaling Technology, Danvers, Massachusetts), CD79a (clone EPR26537-114, Abcam, Cambridge, England), and SOX10 (clone EPR4007-104, Abcam, Cambridge, England), each paired with its respective species-specific HRP polymer (MACH2 Rabbit-HRP for all except FoxP3, above, and SOX10, which used MACH4 Universal HRP) and Opal fluorophore (Opal 520, 620, 480, 690, 570, and TSA-DIG, respectively; Akoya Biosciences, Marlborough, Massachusetts). Microwave retrieval was performed in AR6 after rounds 1–3 and in AR9 after rounds 4–6.

For the myeloid panel (Fig. 4c), primary antibodies were applied in the order neutrophil elastase (clone E8U3X, Cell Signaling Technology, Danvers, Massachusetts; Da Vinci Green Diluent, 30 minutes, room temperature), Ly6G + Ly6C (clone RB6-8C5, Abcam, Cambridge, England; Rat 1-Step HRP, 10 minutes polymer; Biocare, Concord, California), CD11b/ITGAM (clone E4K8C, Cell Signaling Technology, Danvers, Massachusetts), CD68 (clone E3O7V, Cell Signaling Technology, Danvers, Massachusetts), PD-L1 (clone F3Z1T, Cell Signaling Technology, Danvers, Massachusetts), and SOX10 (clone EPR4007-104, Abcam, Cambridge, England), each paired with its respective species-specific HRP polymer (MACH2 Rabbit-HRP for all except Ly6G/Ly6C, above, and SOX10, which used MACH4 Universal HRP) and Opal fluorophore (Opal 620, 570, 690, 480, 520, and TSA-DIG, respectively). Microwave retrieval was performed in AR6 after rounds 1–3 and in AR9 after rounds 4–6.

For both panels, a final incubation with Opal 780 (Akoya Biosciences, Marlborough, Massachusetts) diluted in Renoir Red Diluent (Biocare, Concord, California) was carried out for one hour at room temperature, followed by nuclear counterstaining with a 1X dilution of Spectral DAPI (Akoya Biosciences, Marlborough, Massachusetts). Slides were then removed from the autostainer and coverslipped with ProLong Diamond antifade mountant (Fisher Scientific, Pittsburgh, Pennsylvania). Reagents used for mpIF are summarized in Table S5.

Slides were imaged using the Vectra Polaris Automated Quantitative Pathology Imaging System (Akoya Biosciences, Marlborough, Massachusetts), capturing 1 mm core images for analysis using inForm image analysis software (Akoya Biosciences, Marlborough, Massachusetts). A base algorithm was developed to segment tissue based on SOX10, DAPI, and autofluorescence signal into epithelium, stroma, and other regions (e.g., blank space, tissue folds, and areas of advanced necrosis). Individual phenotyping algorithms were developed to identify mutually exclusive marker combinations (e.g., CD8–CD20, FoxP3– SOX10, CD68–SOX10, CD3–CD79a). Outputs from inForm were combined in Phenoptr (Akoya Biosciences, Marlborough, Massachusetts) to overlay markers and define composite phenotypes. The finalized algorithms were then applied across the full batch of images, and the resulting data were combined and analysed using PhenoptrReports (Akoya Biosciences, Marlborough, Massachusetts) to generate absolute counts and densities for each cell type in both the epithelial and stromal tissue compartments.

### Statistical analysis

Data analysis was performed using Prism version 11.0 (GraphPad, Boston, Massachusetts) and analysed and presented as stated in individual figure legends. P < 0.05 was considered statistically significant and P values are represented as *P < 0.05, **P < 0.01, ***P < 0.001, and ****P < 0.0001, with specific P values indicated in some figures. All experiments were repeated at least once.

## Acknowledgements

We thank the researchers, animal care and administrative staff who helped make this study possible and the organizations that provided funding: BC Cancer Foundation, National Sciences and Engineering Research Council of Canada (grant #490760-15), Canadian Institutes of Health Research (grant #553608), and Innovakine Therapeutics Incorporated (Victoria, British Columbia, Canada).

## Author contributions

S.C.P., M.E.F., C.P., J.S., M.J.B., and B.H.N. conceived the study and designed receptors and experiments. S.C.P., M.E.F., K.M., K.S., M.A., A.P.L., M.J., S.G., T.B., N.S., N.T., M.Z.X., D.K., and J.S. performed experiments and analyzed data. S.C.P., M.E.F., and B.H.N. wrote the manuscript. All authors reviewed, edited, and approved the final manuscript.

## Materials & Correspondence

Address correspondence requests for materials to: Brad H. Nelson, Deeley Research Centre, BC Cancer, 2410 Lee Avenue, Victoria, British Columbia, V8R 6V5, Canada. Phone: 250.519.5700.

