## Supplemental for "Engineered T cells controlled by granulocyte colony-stimulating factor eradicate murine melanoma without lymphodepletion or interleukin-2"

1 **Supplemental Tables and Figures**

| Construct | Amino acid sequence |
| --- | --- |
| **Human G2R** | ATMARLGNCSTWAAALIILLPGSLEECGHISVSAPIVHLGDPITASCIKQNCSHLDPEPQILWRLG<br>AELQPGGRQQRLSDGTQESIITLPHLNHTQAFLLSCCLNWGNSLQILDQVELRAGYPPAIPHNLSCL<br>MNLTTSSLICQWEPGPETHLPTSFTLKSFKSRGNCQTQGDSILDCVPKDGQSHCCIPRKHLLLYQN<br>MGIWVQAENALGTSMSPLCLDPMDEVKLEPPMLRTMDPSPEAAPPQAGCLQLCWEPWQPG<br>LHINQKCELRHKPQRGEASWALVGPLPLEALQYELCGLLPATAYTLQIRCIRWPLPGHWSWDWSPSL<br>ELRTTERAPTVRDLTWWRQRQLDPRTVQLFWKVPLEEDSGRIQGYVVSWRPSGQAGAILPLCN<br>TTELSCTFHLPSAEQVALVAYNSAGTSRPTPVVFSESRGPALTRLHAMARDPHSLWVGWEPNP<br>WPQGYVIEWGLGPPSASNSNKTWRMEQNGRATGFLLENIRPFQLYEIIIVTPLYQDTMGPSQHV<br>YAYSQEMAPSHAPELHLKHIGKTWAQLEWVPEPPELGKSPLETHYTFWTNAQNQSFASILNASSR<br>GFLVHGLEPASLYHIHLMAASQAGATNSTVLTMTLTPEGSELHIILGLFGLLLLTCLCGTAWLCCS<br>PNRKNPLWPSVDPAPHSSLSWVPTIMEEDAFQLPGLGTPPITKLTVLEDEKKPVPWESHSSNH<br>SLTSCFTNQGYFFHLPDALEIEACQVYFTYDPYSEEDPDEGVAGAPTGSSPQLQLPSGEDDAYCT<br>FPSRDDLLLFSPSLGGPSPSTAPGGSGAGEERMPPSLQERVPRDWDQPPLGPPTPGVPDLVDF<br>QPPPELVLRAGEEVPDAGPREGVSFPWSRPPGQGEFRALNARLPLNTDAYLSLQELQGQDPTHL<br>V |
| **Mouse G2R** | ATMVGLGACTLTGVTLIFLLPRSLESCGHIEISPPVVRGDPVLASCTISPNCCKLDQQAQILWRLQ<br>DEPIQPGDRQHHLDPDGTQESLITLPHLNHTQAFLLFCLVPWEDSVQLLDQAEHAGYPPASPSNLSC<br>LMHLTTNSLVCQWEPGPETHLPTSFILKSFRRADQCQYQGDITPDCVAKKRQNNCSIPRKNLLLYQ<br>YMAIWVQAENMLGSSESPKLCCLDPMDEVKLEPPMLQALDIGPDVVSHPGCLWLSWKPKPS<br>EYMEQECELRYQPQLKGANWTLVFHLPSSKDQFELCGLHQAQVYTLQMRIRSSLPGFWSWSP<br>GLQLRPTMKAPTIRLDTCQKKQLDPGTVSQVQLFWKPTPLQEDSGQIQGYLLSWNSPDHQQD<br>IHLCNTTQLSCIFLLPSEAQNVTLVAYNKAGTSSPTTVFLENEGPAVTGLHAMAQDLNTIWDW<br>EAPSLPQGYLIEWEMSSPSYNSYKSWMIENGNITGILLKDNINPFQLYRITVAPLYPGIVGPPV<br>NVYTFAGERAPPHAPALHLKHVGTWTAQLEWVPEAPRLGMIPLTHYTFWADAGDHSFVTLNI<br>SLHDFVLKHLEPASLYHVYLMATSRAGSTNSTGLTLRLDPSDLNIFLGILCLVLLSTTCVVTWLCKK<br>RRGKTSFWSDVPDPAHSSLSWLPTIMTEETFQLPSFWDSSVPSITKITELEEDKKPTHWDSESPS<br>GHSQASCFTNQGYFFFHLPNALEIESCQVYFTYDPCVEEEVEEDGSRLPEGSPHPPLLPLAGEQDD<br>YCAFPPRDDLLLFSPSLSTPNTAYGGSRAPEERSPLSLHEGLPSLASRDLMGLQRPLERMPEGDGE<br>GLSANSSGEQASVPEGNLHGQDQDRGQGPILTLNTDAYLSLQELQAQDSVHLI |

2 **Table S1:** Protein sequences for human and mouse G2R.

| **Pmel-1**<br>**T cells** | **Cytokine** | **Study 1** | **Study 1 re-challenge** | **Study 2** | **Study 2 re-challenge** | **Study 3** | **Study 3 re-challenge** |
| --- | --- | --- | --- | --- | --- | --- | --- |
| N/A (PBS) | Vehicle (3 x weekly) | N/A | N/A | 0/5 CR | N/A | N/A | N/A |
| Mock | Vehicle (3 x weekly) | 0/4 CR | N/A | 0/8 CR | N/A | N/A | N/A |
| Mock | IL-2 (30KIU), daily | 0/5 CR | N/A | 1/8 CR | 0/1 reject (re-treated with IL-2) | N/A | N/A |
| Mock | PEG-GCSF (3.5µg, 3x weekly) | 0/5 CR | N/A | 0/8 CR | N/A | N/A | N/A |
| Mock | IL-2 (30KIU), daily, + PEG-GCSF (3.5µg, 3x weekly) | N/A | N/A | 0/5 CR | N/A | N/A | N/A |
| G2R | Vehicle (3 x weekly) | 0/5 CR | N/A | 0/8 CR | N/A | 0/5 CR | N/A |
| G2R | IL-2 (30KIU), daily | 0/3 CR | N/A | N/A | N/A | N/A | N/A |
| G2R | PEG-GCSF (3.5µg, 3x weekly) | 4/5 CR | 2/4 reject without G-CSF re-treatment<br>1/2 CR with G-CSF re-treatment<br>Total: 3/4 survivors | 7/8 CR | 5/7 reject without G-CSF re-treatment<br>1/2 CR with G-CSF re-treatment<br>Total: 6/7 survivors | 3/5 CR | 3/3 reject without G-CSF re-treatment<br>Total: 3/3 survivors |

3 **Table S2:** Treatment groups and survival outcomes across three independent studies.

| Target | Clone | Fluorophore | Supplier | Catalogue No. | Volume (μL) used per sample |
| --- | --- | --- | --- | --- | --- |
| CD45 | HI30 | APC/FIRE810 | BioLegend, San Diego, California | 304076 | 0.25 |
| CD3 | SK7 | BV750 | BioLegend | 344846 | 0.50 |
| CD4 | SK3 | AF700 | BioLegend | 344622 | 0.50 |
| CD8 | SK1 | PerCP | BioLegend | 344708 | 0.50 |
| G-CSFR | LMM741 | PE | BioLegend | 346106 | 1.00 |

4 **Table S3:** Antibodies used for flow cytometric detection of surface markers on human T cells.

| Target | Clone | Fluorophore | Supplier | Catalogue No. | Volume (μL) used per sample |
| --- | --- | --- | --- | --- | --- |
| CD45 | 30-F11 | APC/Fire 810 | BioLegend, San Diego, California | 103174 | 0.25 |
| CD3 | 17A2 | BV750 | BioLegend | 100249 | 0.50 |
| CD4 | RM4-5 | PE/Cyanine7 | BioLegend | 100528 | 0.50 |
| CD8 | 53-6.7 | PerCP | BioLegend | 100732 | 0.50 |
| CD90.1 | OX-7 | BV421 | BD Biosciences, Franklin Lakes, New Jersey | 563770 | 0.25 |
| Ly-6G | 1A8 | BV570 | BioLegend | 127629 | 1.00 |
| Ly-6C | HK1.4 | FITC | BioLegend | 128006 | 0.50 |
| CD11b | M1/70 | BV605 | BioLegend | 101257 | 1.00 |
| CD11c | N418 | BV711 | BioLegend | 117349 | 1.00 |
| CD62L | MEL-14 | BV510 | BioLegend | 104441 | 1.00 |
| CD44 | IM7 | BV785 | BioLegend | 103059 | 0.50 |
| PD-1 | 29F.1A12 | PE/Dazzle 594 | BioLegend | 135228 | 1.00 |
| CD19 | 6D5 | Spark NIR 685 | BioLegend | 115568 | 0.50 |
| NK1.1 | PK136 | BV480 | BD Biosciences | 746265 | 0.50 |
| G-CSFR | 723806 | PE | Bio-Techne, Minneapolis, Minnesota | D223113 | 1.00 |

5 **Table S4:** Antibodies used for flow cytometric detection of surface markers on mouse immune
6 cells in peripheral blood samples.

| Reagent (clone number) | Supplier (manufacturer) | Catalogue No. |
| --- | --- | --- |
| Xylene | Fisher Scientific, Pittsburgh, Pennsylvania | HC700 1GAL |
| Reagent Alcohol | Fisher Scientific | HC600 1GAL |
| 10% neutral buffered formalin | Sigma-Aldrich, St. Louis, Missouri | HT501128-4L |
| Diva Decloaker | Intermedico (Biocare, Concord, California) | BC-DV2004 |
| Peroxidase-1 | Intermedico | BC-PX968 |
| Background Sniper | Intermedico | BC-BS966 |
| DaVinci Green Diluent | Intermedico | BC-PD900 |
| Renoir Red Diluent | Intermedico | BC-PD904 |
| Mach2 Rabbit-HRP Polymer | Intermedico | BC-RHRP520 |
| MACH4 Universal HRP Polymer | Intermedico | M4U534 G |
| Automation Wash Buffer TBS, 20X | Intermedico | BC-TWB945 |
| Automation Tween-20, 20X | Intermedico | BC-TWA20M |
| OPAL 480 | Akoya Biosciences, Marlborough, Massachusetts | FP1500001KT |
| OPAL 520 | Akoya Biosciences | FP1487001KT |
| OPAL 570 | Akoya Biosciences | FP1488001KT |
| OPAL 620 | Akoya Biosciences | FP1495001KT |
| OPAL 690 | Akoya Biosciences | FP1497001KT |
| OPAL 780 Reagent Pack | Akoya Biosciences | FP1501001KT |
| 1X amplification diluent | Akoya Biosciences | FP1498 |
| AR6 | Akoya Biosciences | AR600250ML |
| AR9 | Akoya Biosciences | AR900250ML |
| DAPI | Akoya Biosciences | FP1490 |
| Prolong Diamond Antifade | Thermo Fisher Scientific, Waltham, Massachusetts | P36961 |
| CD3 (SP7) | Abcam, Cambridge, England | ab16669 |
| FoxP3 (FJK-16s) | Thermo Fisher Scientific | v14-5773 |
| CD20 (E3N7O) | Cell Signaling Technology, Danvers, Massachusetts | 70168 |
| CD8a (D4W2Z) | Cell Signaling Technology | 98941 |
| CD79a (EPR26537-114) | Abcam | ab300150 |
| SOX10 (EPR4007-104) | Abcam | ab180862 |
| Neutrophil Elastase (E8U3X) | Cell Signaling Technology | 90120 |
| Ly6C/Ly6G (RB6-8C5) | Abcam | ab25377 |
| CD11b/ITGAM (E4K8C) | Cell Signaling Technology | 93169 |
| CD68 (E3O7V) | Cell Signaling Technology | 97778 |
| PD-L1 (F3Z1T) | Cell Signaling Technology | 80517 |

7 **Table S5:** Reagents and antibodies used for mpIF staining.

**Fig. S1**

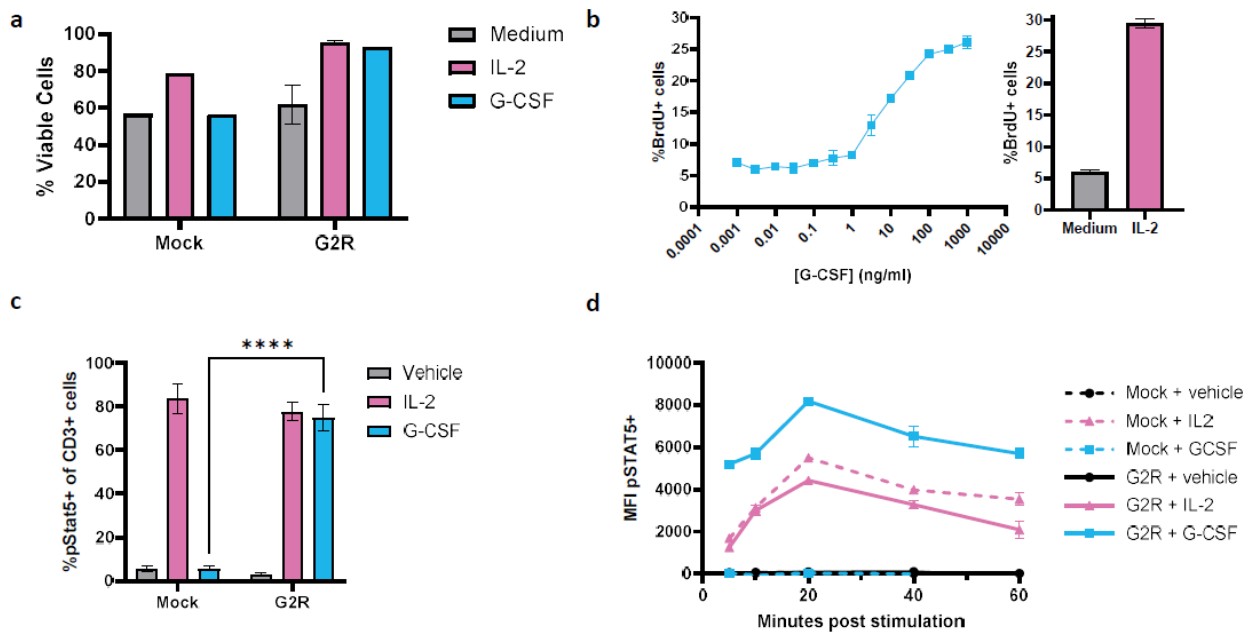

**Supplemental Figure 1. Viability, proliferation and STAT5 phosphorylation in G2R-expressing**

**human T cells stimulated with G-CSF versus IL-2.** Human T cells expressing G2R or mock-

transduced T cells were assessed in vitro for cytokine responsiveness. **(a)** Viability of T cells on

day 20 of expansion. **(b)** Incorporation of BrdU by G2R-expressing T cells (blue) as a function of

increasing concentration of G-CSF, with comparison to medium alone (grey) or IL-2 (pink). **(c-d)**

T cells expressing G2R or mock-transduced were stimulated with vehicle, IL-2 or G-CSF and

analysed for STAT5 tyrosine phosphorylation by flow cytometry. **(c)** Phospho-STAT5 positivity in

T cells in response to a 20-minute stimulation with vehicle, IL-2, or G-CSF. Unpaired t test with

Welch's correction. \*\*\*\*P < 0.0001. **(d)** MFI of phospho-STAT5-phycoerythrin (PE) in T cells in

response to a stimulation with vehicle, IL-2, or G-CSF at the indicated timepoints.

Fig. S2

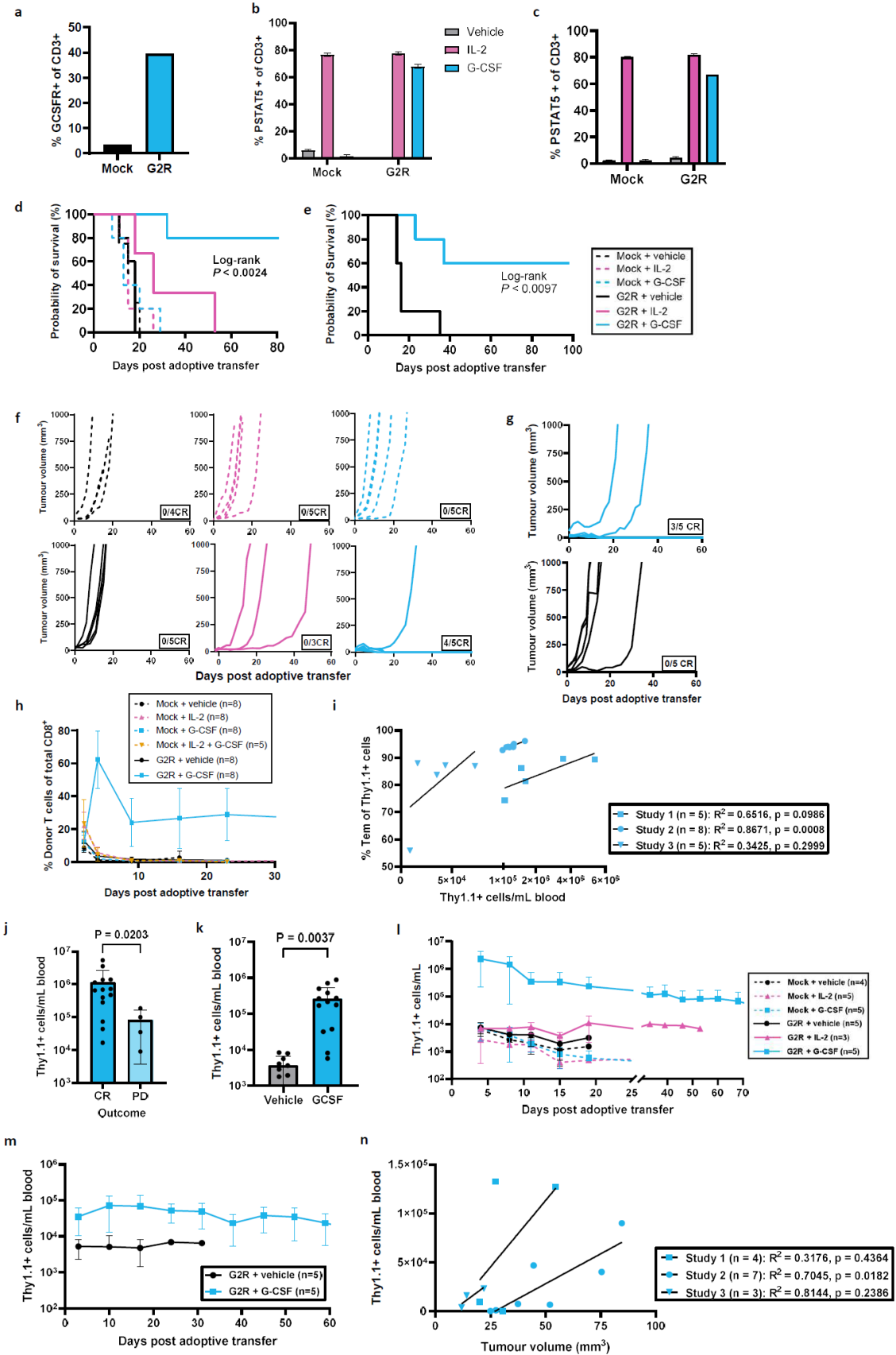

**Supplemental Figure 2. Additional data demonstrating that G-CSF drives expansion of G2R-expressing pmel-1 T cells and tumour regression in B16-F10 tumour-bearing mice.** C57BL/6J mice (n = 3 to 8 per group, as per Supplemental Table S2) were subcutaneously implanted with 5 x 10<sup>5</sup> B16-F10 cells seven days prior to intravenous adoptive transfer of 1 x 10<sup>7</sup> pmel-1 T cells expressing G2R or mock-transduced. Mice were randomized to receive intraperitoneal injections of G-CSF, IL-2 or vehicle control. **(a)** G2R surface expression on pmel-1 T cells used for ACT in Study 3, as measured by expression of G-CSFR by flow cytometry. **(b-c)** Phospho-STAT-5 expression in **(b)** Study 2 and **(c)** Study 1, in G2R- or mock-transduced pmel-1 T cells, in response to a 20-minute stimulation with vehicle, IL-2 (300 IU/mL) or PEG-GCSF (100 ng/mL). **(d-e)** Kaplan-Meier survival curves for **(d)** Study 1 and **(e)** Study 3, each compared by log-rank (Mantel-Cox) test. **(f-g)** Tumour volumes as measured by caliper three times weekly for **(f)** Study 1 and **(g)** Study 3. **(h)** Adoptively transferred pmel-1 T cells in circulation, as a percentage of total circulating CD8<sup>+</sup> T cells, measured by flow cytometry of peripheral blood samples from Study 2. **(i)** Correlation between Thy1.1<sup>+</sup> pmel-1 T cells and %Tem on day 3 (Study 3) or day 4 (Studies 1 and 2) post-ACT in mice treated with G2R<sup>+</sup> pmel-1 T cells + G-CSF. Correlation was assessed using a two-tailed Pearson correlation test. The solid line indicates the linear regression fit. Each point represents an individual mouse. **(j)** Circulating Thy1.1<sup>+</sup> pmel-1 T cells on day 3 or 4 (depending on the study) and survival outcome (complete regression [CR] vs. progressive disease [PD]) of mice in G2R<sup>+</sup> + G-CSF treatment group across Studies 1 to 3. Unpaired two-tailed t test with Welch's correction used to compare means. **(k)** Circulating Thy1.1<sup>+</sup> pmel-1 T cells on day 15, 16, or 17 (depending on the study) in mice that were treated with G2R<sup>+</sup> T cells + vehicle versus G2R<sup>+</sup> T cells + G-CSF and exhibited CR. Data is compiled across Studies 1 to 3. Unpaired two-tailed t test with Welch's correction used to compare means. **(l-m)** Circulating Thy1.1<sup>+</sup> pmel-1 T cells over time in **(l)** Study 1 and **(m)** Study 3. **(n)** Correlation between tumour volume on day 1 (Study 1) or 2 (Studies 2 and 3) post-ACT and Thy1.1<sup>+</sup> pmel-1 T cells on day 66 (Study 3), 68 (Study 1), or 70 (Study 2) post-ACT in mice treated with G2R<sup>+</sup> pmel-1 T cells + G-CSF. Correlation was assessed using a two-tailed Pearson correlation test. The solid line indicates the linear regression fit. Each point represents an individual mouse.

**Fig. S3**

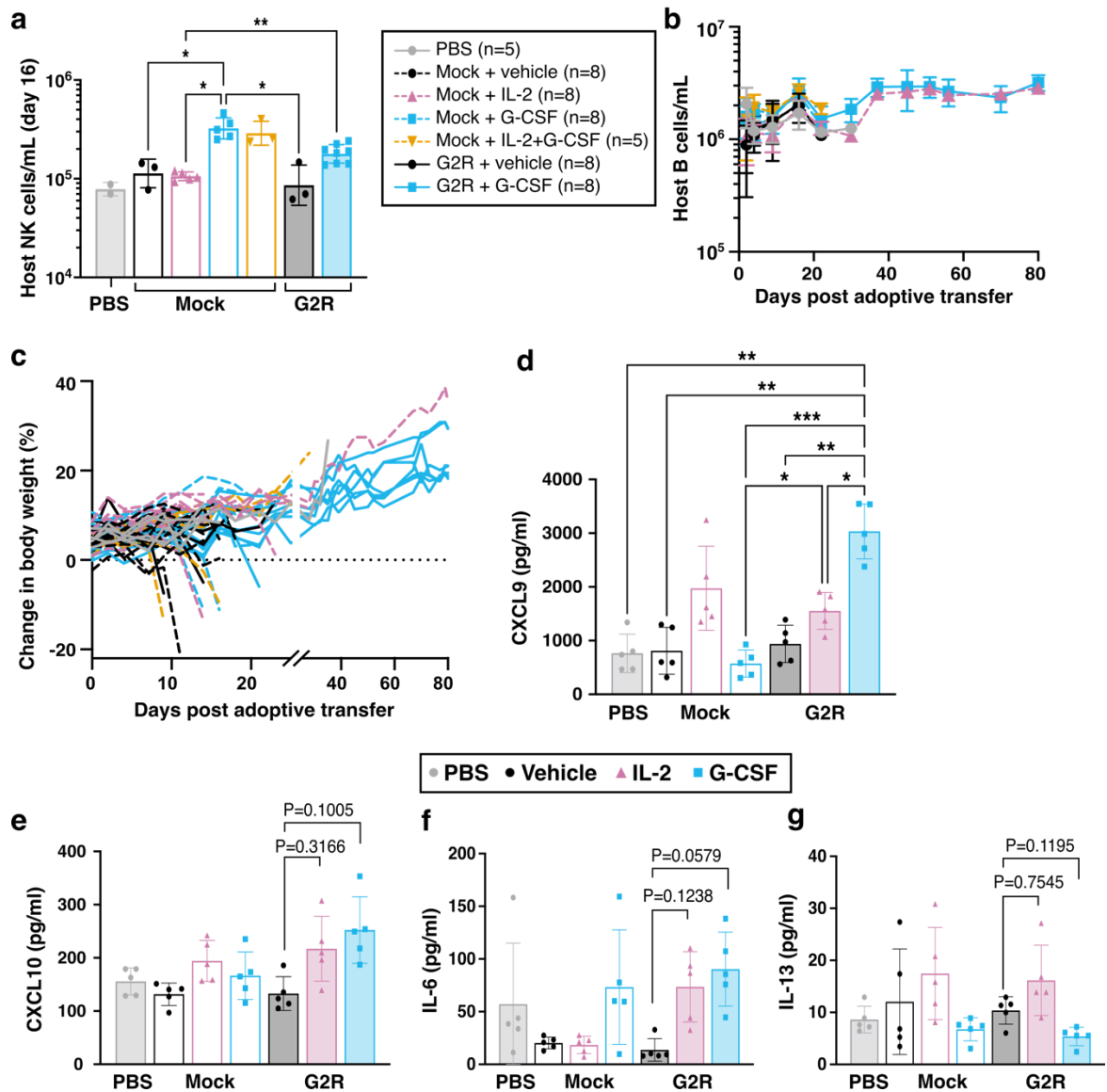

**Supplemental Figure 3. Host immune and systemic responses to G-CSF-driven expansion of G2R-expressing T cells in B16-F10-bearing mice.** C57BL/6J mice (n = 5-8 per group, as per Supplementary Table S2) were subcutaneously implanted with $5 \times 10^5$ B16-F10 cells seven days prior to intravenous adoptive transfer of $1 \times 10^7$ pmel-1 T cells expressing G2R or mock-transduced. Mice were randomized to receive intraperitoneal injections of G-CSF, IL-2 or vehicle control. **(a)** Circulating NK cells on day 16 post adoptive transfer in Study 2. **(b)** Circulating B cells on day 16 post adoptive transfer in Study 2. **(c)** Percentage change in body weight since time of tumour implant of mice across all treatment groups in Study 2. **(d-g)** Cytokine levels in plasma collected on day 4 after adoptive transfer of G2R<sup>+</sup> T cells or mock-transduced T cells followed by treatment with vehicle, IL-2 or G-CSF as indicated. Data presented as mean of five
biological replicates for **(d)** CXCL9, **(e)** CXCL10, **(f)** IL-6, and **(g)** IL-13. Panels **a** and **d-g**: Dunnett's
T3 multiple comparisons test. \*P < 0.05; \*\*P < 0.01; \*\*\*P < 0.001. Data are presented as means
+/- SD of 5 to 8 biological replicates.

Fig. S4

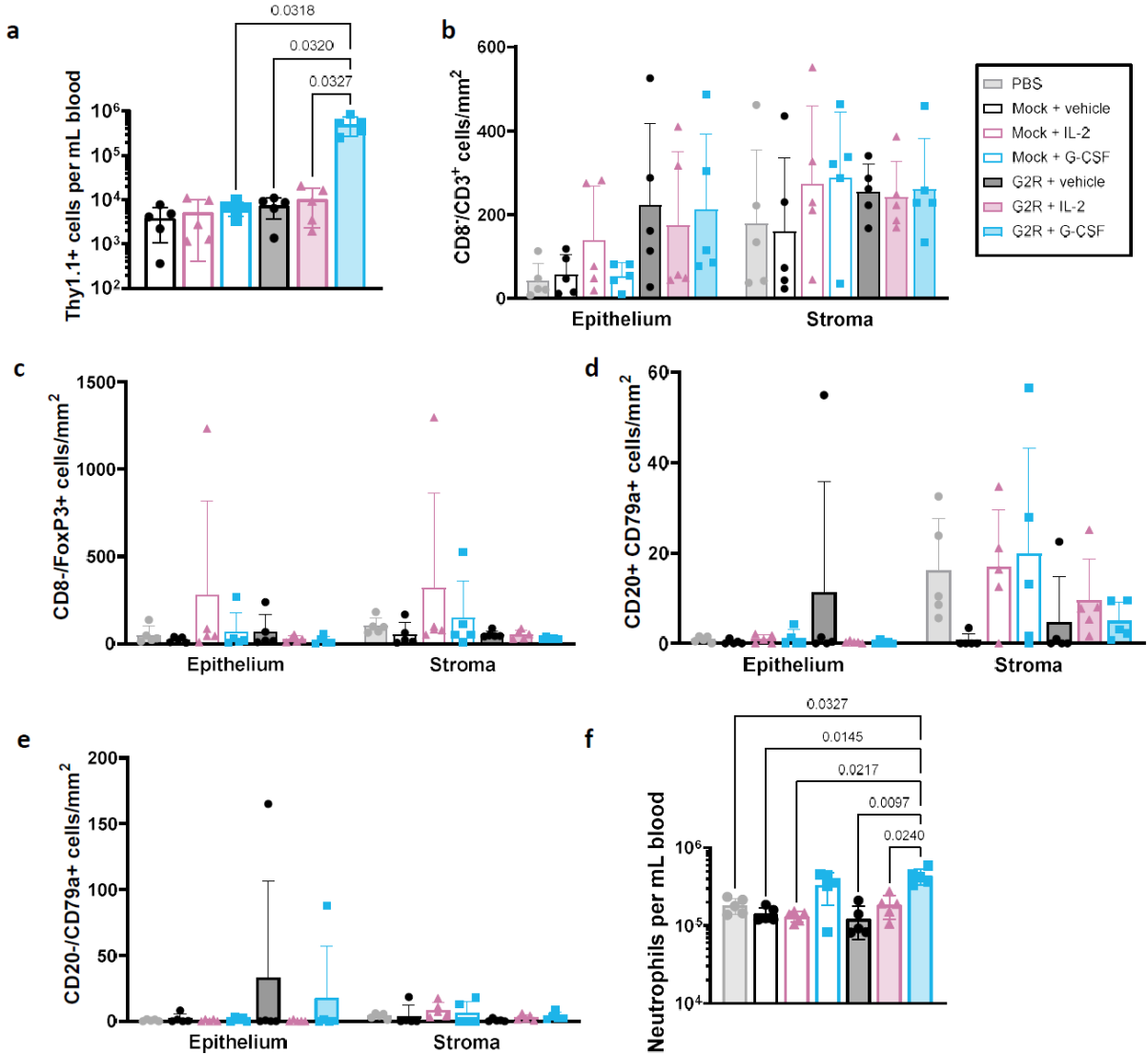

#### Supplemental Figure 4. Day 4 circulating and tumour-infiltrating immune cell profiles

following adoptive transfer of mock-transduced or G2R<sup>+</sup> T cells. Established subcutaneous

B16-F10 melanoma tumours (n = 5 per group) and peripheral blood were collected on day 4

post adoptive transfer of mock-transduced or G2R<sup>+</sup> pmel-1 T cells followed by systemic

administration of G-CSF, IL-2 or vehicle. **(a)** Circulating Thy1.1<sup>+</sup> pmel-1 T cells across treatment

groups, as measured by flow cytometry. **(b-e)** Tumours were analysed by mpIF and the

indicated immune cell subsets were quantified in tumour epithelium and stroma. **(b)**

Presumptive CD4<sup>+</sup> T cells (CD3<sup>+</sup> CD8<sup>-</sup>). **(c)** Presumptive Tregs (CD8<sup>-</sup> FoxP3<sup>+</sup>). **(d)** B cells (CD20<sup>+</sup>

CD79a<sup>+</sup>). **(e)** Plasma cells (CD20<sup>-</sup> CD79a<sup>+</sup>). **(f)** Circulating neutrophils across treatment groups, as measured by flow cytometry. Panels **a** and **f**: One-way ANOVA with Dunnett's T3 multiple
comparisons test. Data are presented as means ± SD of five biological replicates.
